# Large-scale Inference of Population Structure in Presence of Missingness using PCA

**DOI:** 10.1101/2020.04.29.067496

**Authors:** Jonas Meisner, Siyang Liu, Mingxi Huang, Anders Albrechtsen

## Abstract

**Background:** Principal component analysis (PCA) is a commonly used tool in genetics to capture and visualize population structure. Due to technological advances in sequencing, such as the widely used non-invasive prenatal test, massive datasets of ultra-low coverage sequencing are being generated. These datasets are characterized by having a large amount of missing genotype information. We present `EMU`, a method for inferring population structure in the presence of rampant non-random missingness.

**Results:** We show through simulations that several commonly used PCA methods can not handle missing data arisen from various sources, which leads to biased results as individuals are projected into the PC space based on their amount of missingness. In terms of accuracy, `EMU` outperforms an existing method that also accommodates missingness while being competitively fast. We further tested `EMU` on around 100K individuals of the Phase 1 dataset of the Chinese Millionome Project, that were shallowly sequenced to around 0.08x. From this data we are able to capture the population structure of the Han Chinese and to reproduce previous analysis in a matter of CPU hours instead of CPU years.

**Conclusions:** `EMU`’s capability to accurately infer population structure in the presence of missingness will be of increasing importance with the rising number of large-scale genetic datasets. `EMU` is written in `Python` and is freely available at https://github.com/Rosemeis/emu/.

## 1 Introduction

The advent of whole-genome sequencing technologies has brought the opportunity of generating large amount of genomic data at low cost [25]. Large-scale sequencing studies are therefore becoming more prevalent [6, 7, 10, 13, 21] as they help researchers understand genetic variation in populations on a much broader scale than previously possible using genotyping arrays. A cost-effective strategy with the ever-increasing demand for larger sample sizes seems to advocate for the use of medium or low coverage sequencing [26, 20]. Larger sample sizes sequenced at lower depths will generally lead to better population-scale estimates of genetic variation compared to sequencing at higher depths at the cost of limited sample sizes [11]. With this appealing trade-off, a genomic study [21] was recently conducted on ultra-low coverage sequencing data of 141K Chinese pregnant women as part of the Chinese Millionome Project. The participants in the study underwent a non-invasive prenatal test (NIPT) which is common for testing fetal chromosomal abnormalities by shallowly sequencing the cell-free DNA from the maternal plasma. The study provided insight into the genetic structure and history of the Chinese population as well as performing genome-wide association studies (GWAS) with principal components as covariates. The study had an average depth of < 0.1X, which allowed for the much larger sample size compared to other sequencing projects.

These large-scale sequencing studies will usually consist of individuals from diverse ancestries and may include cryptic structure not accounted for. Population structure plays a major role in population genetics for understanding population demography [27], as well as in association studies where it acts as a confounding factor and must be accounted for [22, 29]. One approach for inferring population structure is based on the use of principal component analysis (PCA). PCA has the appealing feature of projecting individuals onto inferred axes of genetic variation that capture population structure in a continuous fashion. The standard way to infer population structure using PCA has been to construct a genetic relationship matrix (GRM) and perform eigendecomposition on this matrix to infer the axes of genetic variation [27]. However as the sample size in largescale studies is constantly increasing, faster and more scalable methods based on various low-rank approximations [19, 14] have been developed to only infer the top axes of genetic variation almost directly from the genotype matrix [23, 12, 1]. The problem for the majority of these methods is that they cannot handle missing data in an appropriate manner.

Common approaches for dealing with missingness when inferring population structure are either to thin the dataset by removing sites or individuals with missingness rates above a certain threshold or to simply ignore the presence of missingness by using the mean genotype value [27, 12, 1], also called mean imputation. The problem with the first approach is that one would lose information that could potentially be crucial in downstream analyses, and especially when merging datasets of different sources. The problem with second approach is that the inferred axes will correlate with the amount of missingness and thus no longer only represent population structure. This is due to missingness being modelled as the average across the entire dataset.

We therefore propose a new method that is specially designed to deal with large-scale genetic datasets with very high levels of missingness using a novel accelerated approach. `EMU` (EM-PCA for Ultra-low Coverage Sequencing Data) is an accelerated expectation-maximization (EM) algorithm for PCA to model the missingness in an iterative fashion. The concept of an iterative PCA for imputing missing values is not novel and has been formally described [17, 16]. A similar method [23] has also been developed for low coverage sequencing data on the basis of genotype likelihoods which however is not optimal for ultra-low coverage sequencing data, where individuals almost never have more than one read at a locus. We apply and show that our method displays high accuracy, robustness and scalability on both simulated and real data with very high missingness, where we apply it to around 100K non-invasive prenatal test data [21]. In relation to missingness patterns, we also demonstrate that `EMU` is robust to having different SNP ascertainment schemes in a dataset as would be a result of merging different data types. Additionally, we compare our method in terms of accuracy, computational speed and memory usage against other popular choices of methods for inferring population structure on the basis of PCA.

## 2 Material and Methods

We will now describe our method `EMU` for inferring population structure in ultra-low coverage sequencing data. As we assume to be working on datasets with extensive amount of missingness for ultra-low coverage sequencing data (≤ 0.1X), we use a single-read sampling approach to best describe our data in a similar fashion to [21]. This means that we expect on average ≤ 0.1 sequencing reads to map to a given position in the covered genome on average.

We therefore define a data matrix **D** with its entries representing the output of a single-read sampling approach for observed sequencing reads in *n* individuals and *m* variable sites. Thus for *i* = 1, …, *n* and *j* = 1, …, *m, d*_*ij*_ can take values from {0, 1, −9} where 0 and 1 are the sampling of the major and minor allele, respectively, while −9 refers to missing data for the given individual *i* in the given site *j*. We thereby assume that all sites are diallelic and that both major and minor alleles are known and we ignore sequencing errors. The sampling of an allele can therefore be seen as a Bernoulli process. `EMU` is also capable of working with diploid genotype data, where *d*_*ij*_ can take values from {0, 0.5, 1, −9} such that heterozygous genotype information is kept, but here we describe it for pseudo-haploid data (single-read sampling).

### 2.1 Population allele frequencies

The allele frequency across all samples in the dataset 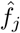 is estimated as follows for a single site *j* by counting the observed number of minor alleles. We denote this as the population allele frequency:

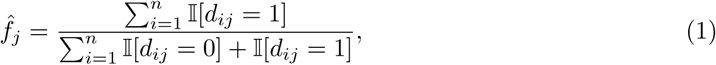

with 𝕀[*X*] being the indicator function. I.e. the allele frequencies are calculated such that individuals with missing information are not counted for the given site. For diploid genotype data, we also count the heterozygous individuals (*d*_*ij*_ = 0.5) in both numerator and denominator.

### 2.2 Individual allele frequencies

Pritchard et al. [30] introduced the concept of individual allele frequencies in `STRUCTURE` for genotype data. Under the assumption of a fixed number of ancestral populations, admixture proportions **Q** and ancestral allele frequencies **F** are inferred that when multiplied represent individual allele frequencies, **П** = **QF**^*T*^. More recently Hao et al. [15] constructed a similar approach for genotype data, instead based on PCA, where the top principal components are used to reconstruct the genotype matrix, and the individual allele frequencies can be derived from a low-rank approximation. Several methods have applied this idea [23, 5, 24]. In this study, we present an iterative variant for pseudo-haploid data that accounts for missingness.

For individual *i* at site *j* the individual allele frequency *π*_*ij*_ can be seen as the underlying parameter in the binomial sampling process of genotype *g*_*ij*_, conditioned on the population structure.

### 2.3 Iterative PCA

`EMU` is based on the iterative PCA algorithm of [17] (EM-PCA) which deals with finding a low-rank approximation of **D** iteratively, where missing values are imputed by reconstruction from the estimated low-rank approximation of the previous iteration. This iterative procedure corresponds to an EM algorithm and it is equivalent to finding the matrix of individual allele frequencies that minimize the expression [16]

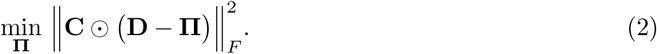

Here **C** is a weight matrix with entries such that *c*_*ij*_ = 0 if *d*_*ij*_ = −9 and *c*_*ij*_ = 1 otherwise for *i* = 1, …, *n* and *j* = 1, …, *m*, while ⊙ represents element-wise multiplication. Thus, only entries with information are evaluated. However, note that **П** will be estimated from the full dataset. The iterative procedure and its updating scheme to estimate the individual allele frequencies are described in **Algorithm 1**, where missing values are initialized as the population allele frequencies 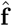 and *K* dimensions are used in the low-rank approximation.

#### Algorithm 1: EM-PCA in EMU

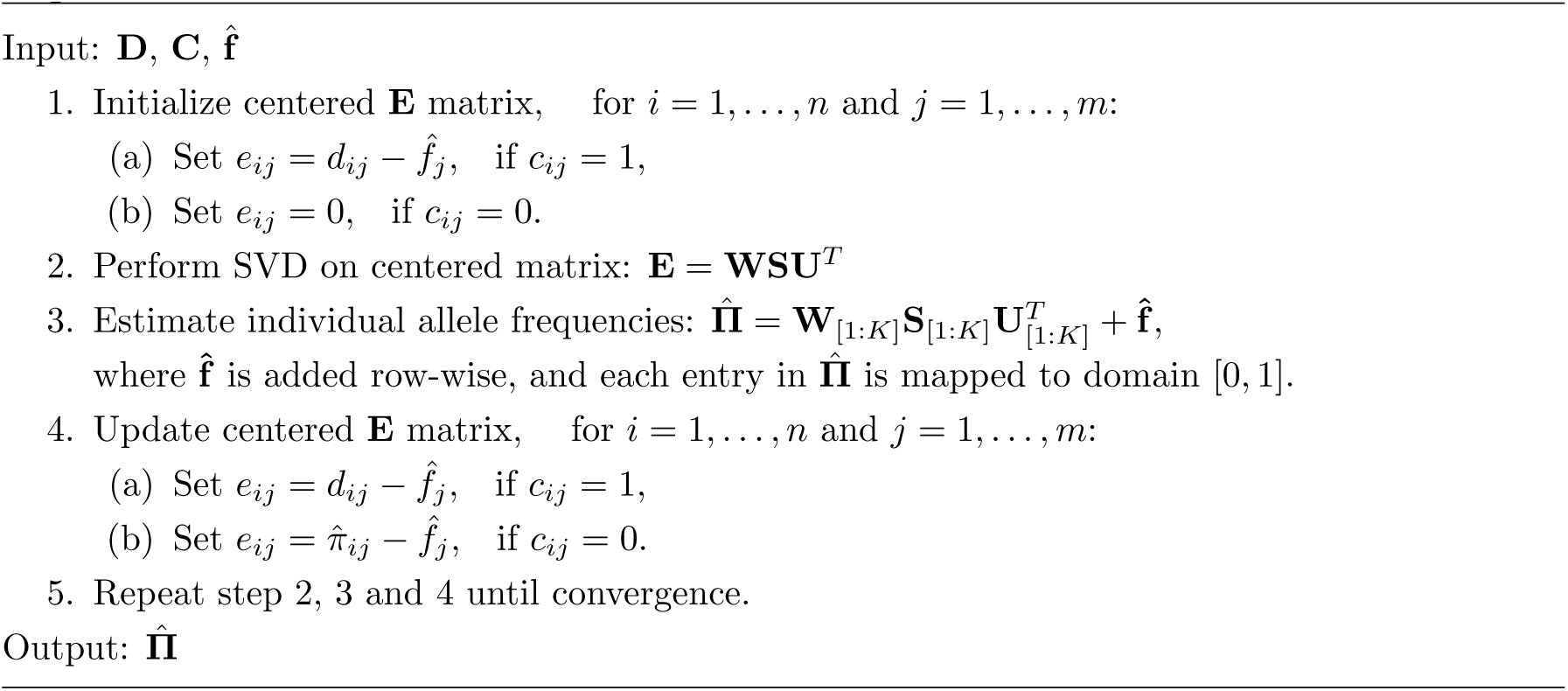

We define convergence as when the root-mean-square deviation (RMSD) of **U**_[1:*K*]_ between two successive iterations of the iterative procedure is less than some small value *ϵ* = 5 × 10^−7^

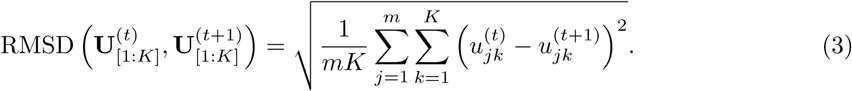

After obtaining the final set of individual allele frequencies that minimizes Equation 2 from our EM-PCA algorithm, we can infer the population structure from a standardized matrix such that the variable sites are weighted by their population allele frequencies, 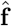 [27]. Thus, define the standardized matrix **X** with entries:

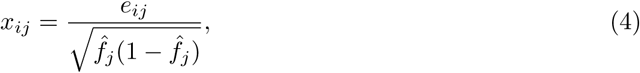

for *i* = 1, …, *n* and *j* = 1, …, *m* with *e*_*ij*_ being defined as step 4 in the final iteration of Algorithm 1. Performing SVD on the standardized matrix will infer and map individuals onto axes of genetic variation that represents population structure (principal components):

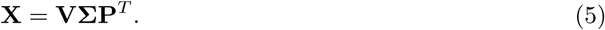

Here **V** will represent the principal components capturing population structure, which are identical to the principal components inferred when performing full eigendecomposition on the genetic relationship matrix [27, 9].

### 2.4 Accelerated EM

As the number of missing values in our data matrix will increase the difficulty of our minimization problem (Equation 2), the parameter space of our algorithm will also increase and it may lead to slow convergence. To speed up the convergence of the EM algorithm, we have implemented an accelerated variant of the iterative update based on the SQUAREM acceleration schemes [32], where we are using SqS3 to update the factor matrices (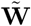 and **U**, with 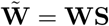) independently. It works by taking two steps using the standard EM algorithm to find an optimal larger step in the parameter space based on a combination between the previous and new estimates (step 2, 3 and 4 in Algorithm 1).

### 2.5 Implementation

`EMU` has been implemented in `Python` using `Numpy` [31] data structures and `Cython` [2] for parallelization and to speed up computational bottlenecks. We are using a truncated SVD implementation of the `scikit-learn` library [28] that uses a randomized PCA procedure [14] to only compute the *K* largest eigenvectors of a given matrix. We have also extended our method to work on diploid genotypes such that the information of heterozygous sites are retained.

The code is freely available at https://github.com/Rosemeis/emu/ and works using both `Python` 2.7 and 3.7.

The computational complexity for one iteration in our EM-PCA algorithm will be 𝒪(*nm*) for low-rank approximations using the truncated SVD procedure, and the number of iterations will depend on the amount of missingness and number of samples and variable sites in the dataset. In total, our algorithm will have a memory requirement of 𝒪(*nm*) bytes. However in modern largescale datasets, the constant to this bound will be important for actual applications. A more detailed description of the memory usage is described in the supplementary material. The algorithm is linear in both the number of samples and the number of sites for both computational speed and memory usage.

We have also implemented an alternative variant of the iterative update in our algorithm that is much more memory efficient by slightly sacrificing computational speed. This variant uses ∼20x less memory than the previously described procedure by using a lazy evaluation approach for **E** in custom matrix multiplications, based on the same randomized SVD procedure, such that only the low-rank factor matrices of **E** are considered. A description of this variant can also be found in the supplementary material and we will regard to this variant as `EMU-mem` in the main results.

### 2.6 Simulation of single-read data matrix

We have simulated genotype data to test the capabilities of our method. The simulations are based on allele frequencies from three Chinese populations (Han, Uygur and Dai) generated from genotyped individuals [18] of the Human Genome Diversity Project [3]. These populations were selected to make the scenario somewhat similar to the real data used in this study with assumed low *F*_*ST*_ distances between populations. We simulate individuals under a Binomial model, such that a sampled genotype *g*_*ij*_ can take values of 0, 1, and 2, representing the minor allele count for individual *i* at site *j*. We then transform the simulated genotype matrix into a single-read sampling matrix (pseudo-haploid) using the following scheme for individual *i* at site *j*. A homozygous genotype (*g*_*ij*_ ∈ {0, 2}) is directly converted to either 0 or 1, respectively, while a heterozygous genotype (*g*_*ij*_ = 1) is simply converted to either 0 or 1 with equal probability in the single-read sampling matrix. To test and showcase our method in various cases of extreme missingness, we have simulated three different scenarios of missingness as well as five additional scenarios, related to emulating different SNP ascertainment schemes. In all simulated scenarios, the number of variable sites is ∼350K after filtering out rare variants based on a minor allele frequency threshold (5%). To simplify the different scenarios for the reader, we have provided a graphical overview of the simulation procedures in the supplementary material (Figure S1).

#### 2.6.1 Missingness rate scenarios

The three scenarios for different degrees of missingness have been simulated with a total of 900 individuals (250 from each distinct population and 150 being admixed) and a total of 9000 individuals (2500 from each distinct population and 1500 being admixed) to evaluate the effect of sample size.

In Scenario 1, we have simulated individuals with a randomly assigned missingness rate from 5 − 50%. This means that an individual *i* with a missingness rate of 50% for example would have a probability of 0.5 to keep the sampled allele at a given site *j* (*d*_*ij*_ ∈ {0, 1}) and a probability of 0.5 to remove the sampled information (*d*_*ij*_ = −9).

For Scenario 2, we have made the missingness rate gradient more extreme such that the simulated individuals are randomly assigned a missingness rate between 90 − 99%. We have also used this scenario to perform computational tests regarding speed, memory and accuracy for much larger sample sizes later on.

And last in Scenario 3, we have simulated half of the individuals from the three distinct population with a missingness rate sampled from 𝒩(0.95, 0.001) (≈ 95%). The other half and all admixed individuals have been simulated with a missingness rate sampled from 𝒩(0.5, 0.01) (≈ 50%).

#### 2.6.2 SNP ascertainment scheme scenarios

For the five scenarios emulating different SNP ascertainment schemes, we have simulated 750 individuals with 250 from each population such that admixed individuals have not been included.

In Scenario 4, we have simulated a scenario that tries to emulate the merging of a SNP array dataset with a whole-genome sequencing (WGS) dataset. Therefore, we have simulated half of the individuals in each of the three different populations to only retain information of very common variants (MAF ≥ 0.25) and set other variants as missing, while the other halves of the three populations have information from all simulated variants.

For Scenario 5, we have simulated half of the individuals in one of the three different populations (Han) to only retain information in approximately a third of the variants (∼130K), while the other half of the population retain information in the other two-thirds of the variants (∼233K). However, there is a small overlap of 13K between the two subsets. Thus, we create a scenario where one population mostly has to rely on the other two populations to estimate within-population correlations between its two halves. The other two populations (Uygur and Dai) are simulated with full information. The case is similar for Scenario 6, but here two populations (Han and Uygur) have been simulated such that half of their individuals has information in approximately a third of the variants while the other half has information in the other two-thirds of the variants with a slight overlap. The Dai population is simulated with full information.

Scenario 7 and 8 are almost identical to Scenario 5 and 6, however now there is no overlap between the halves of a population. This means that the correlation within a population between its two halves must solely rely on the correlation with the other populations.

The last four scenarios (5, 6, 7 and 8) tries to emulate cases where either different datasets of different SNP arrays have been merged, which may be almost or entirely non-overlapping, or when dealing with ultra-low coverage sequencing data.

### 2.7 Phase 1 of the Chinese Millionome Project

The Chinese Millinome Project aims at analyzing millions of Chinese sequencing genomes to understand the genetic diversity of the Chinese population and to promote the precision medicine initiatives in China (https://db.cngb.org/cmdb/). In the Phase 1 study [21], Liu et al. analysed the low coverage genomes of about 141K female participants that were recruited via the NIPT test during pregnancy. The individuals were sequenced at ultra-low depth with 5-10 million using either 35bp or 49bp single end reads, corresponding to an average sequencing depth of 0.08X. We restricted or analysis to sites that are known to be common (MAF ≥ 0.05) both in our project and in the East Asian populations of the 1000 Genomes Project East Asian population and that had a sequencing depth ≥ 0.1X. We additionally kept individuals sequenced with 35bp reads and removed individuals with sequencing error rate greater than 0.00325[21],. This resulted in pseudo-haploid genotype matrix of 97K individuals and 440K sites.

## 3 Results

For the simulated datasets, we test and compare `EMU` against other commonly used methods for inferring population structure using PCA. These include `PLINK` (version 2.0) [4], `smartpca` and `FastPCA` [12] from the `EIGENSOFT` package (version 7.2.1) [27], `FlashPCA` (version 2.0) [1]. `PLINK` and `smartpca` estimate the GRM and perform full eigendecomposition on this, while `FastPCA` and `FlashPCA` use low-rank approximation approaches on the standardized data matrix. From these methods, `PLINK` is the only other method besides `EMU` that accounts for missingness in the dataset, while the rest perform mean imputation. We also tested multidimensional scaling (MDS) from IBS distances and `SNPRelate` [33] for all simulated scenarios, however we did not include them in the main results. Their results are shown in Figure S16 and S17.

To assess the performance of our method in the various simulated scenarios, we perform Procrustes analyses [8]. We infer population structure from the full genotype datasets, from which the single-read sampling matrices have been generated from, to serve as ground-truth in comparisons with all the tested methods. A Procrustes analysis will then find scaling and rotation components to best represent the inferred principal components of a tested method in comparison with the ground-truth and the RMSD is reported.

### 3.1 Simulation

In the following results, we have tested eight different scenarios of missingness. The first three scenarios are related to individuals having different missingness rates, while the last five scenarios are related to the merging of different datasets and using different SNP ascertainment schemes.

#### 3.1.1 Missingness rate scenarios

There are two cases for each of the three scenario (A and B) and these distinguish the sample size of the simulated dataset. Case A has 900 individuals with 250 individuals sampled from each distinct population and 150 admixed individuals, while case B has 9000 individuals with 2500 individuals sampled from each distinct population and 1500 admixed. We only display the results of case A in the main results while the results of case B are displayed in the supplementary material.

The results of Scenario 1, where the missingness rate is between 5 − 50% for all individuals, are shown in Figure 1 and S2. It can be seen that the three methods, which are not accounting for missingness in the dataset (`smartpca`, `FastPCA` and `FlashPCA`), are struggling to separate the individuals from the three populations into distinct clusters in the presence of missingness and they produce almost identical results where the individuals with more missingness is closer to the origin S5. In contrast, both `EMU` and `PLINK` are able to infer the population structure accurately as also verified in the Procrustes analyses shown in Table 1.

**Table 1:** Procrustes analyses of the methods in the first three simulated scenarios. Measured by RMSD between the PCs inferred from the full dataset (ground-truth) and the transformed PCs inferred from the different tested methods. Scenarios of case A and case B had sample sizes of 900 and 9000, respectively.

| Method | 1A | 1B | 2A | 2B | 3A | 3B |
| --- | --- | --- | --- | --- | --- | --- |
| EMU | **0.000644** | **0.000190** | **0.00390** | **0.00101** | **0.00197** | **0.000556** |
| PLINK | 0.000650 | 0.000192 | 0.00567 | 0.00110 | 0.00262 | 0.000622 |
| FlashPCA | 0.00897 | 0.00266 | 0.0299 | 0.00681 | 0.0292 | 0.00904 |
| smartpca | 0.00897 | 0.00266 | 0.0299 | 0.00681 | 0.0292 | 0.00904 |
| FastPCA | 0.00897 | 0.00266 | 0.0420 | 0.00684 | 0.0292 | 0.00904 |

**Figure 1:**
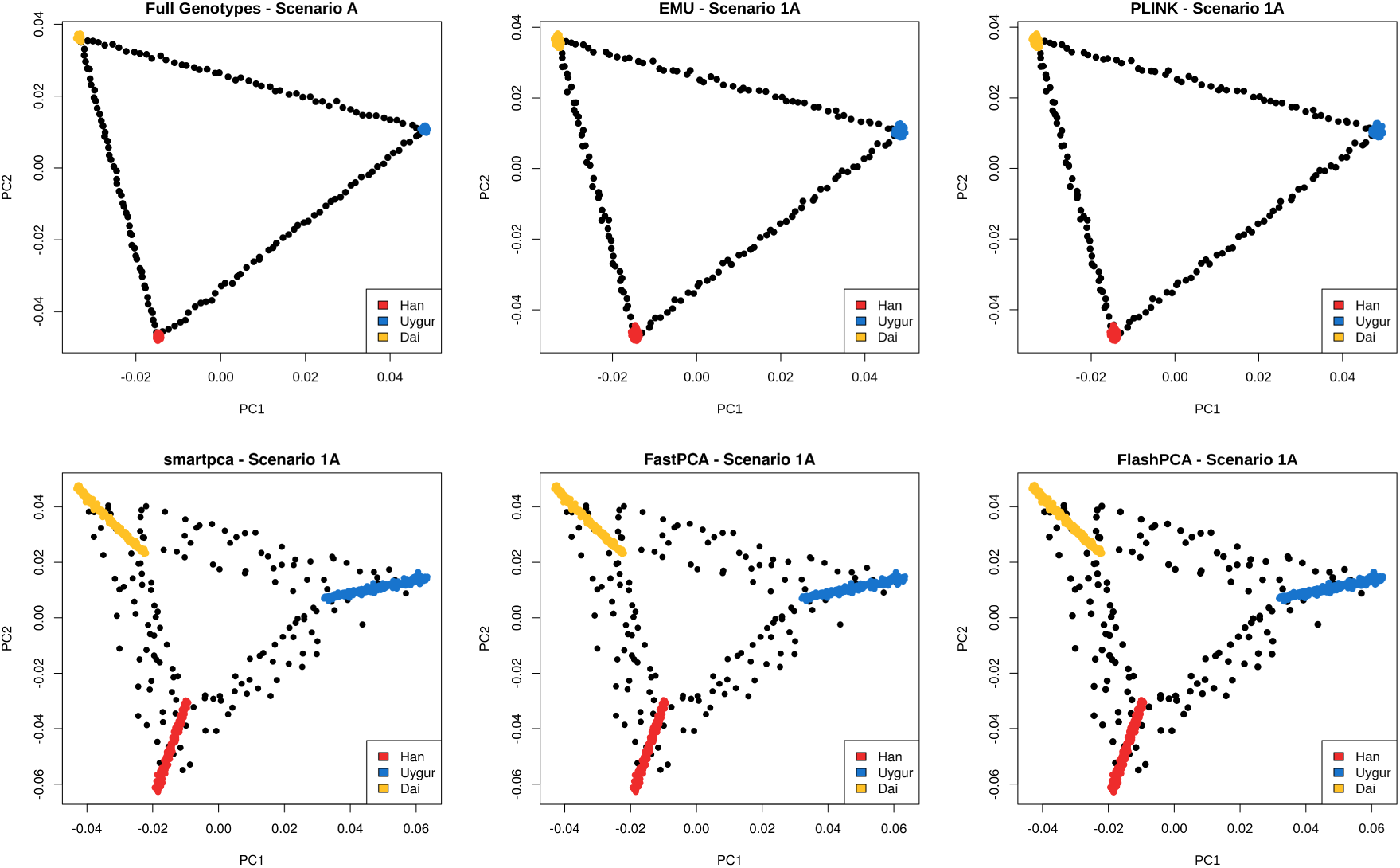
PCA plots of tested methods for Scenario 1A displaying the top two axes of genetic variation for 900 individuals. Black dots represent two-way admixed individuals. The top left plot shows the PCA performed on the full dataset, such that it acts as ground-truth. Individuals were simulated with low to moderate missingness rates between 5 − 50%.

For Scenario 2, the interval of the missingness rate was increased to 90−99% in order to simulate a more extreme scenario as also seen in the Chinese Millionome Project. The results are displayed in Figure 2 and S3 for 900 and 9000 individuals, respectively, and the 900 individuals colored by their missingness rate in Figure S6. Again, `EMU` and `PLINK` are once again able to infer the simulated population structure but `EMU` is able to recover a slightly more accurate PCA plot as shown in Table 1. It can also be seen that the estimates of `PLINK` for the individuals of the three populations are slightly more noisy than the estimates of `EMU`, and that these individuals generally have a higher missingness rate. Here `FastPCA` is not able to infer any meaningful structure in the dataset, while `smartpca` and `FlashPCA` finds some overall pattern but it is heavily biased by the missingness in the dataset.

**Figure 2:**
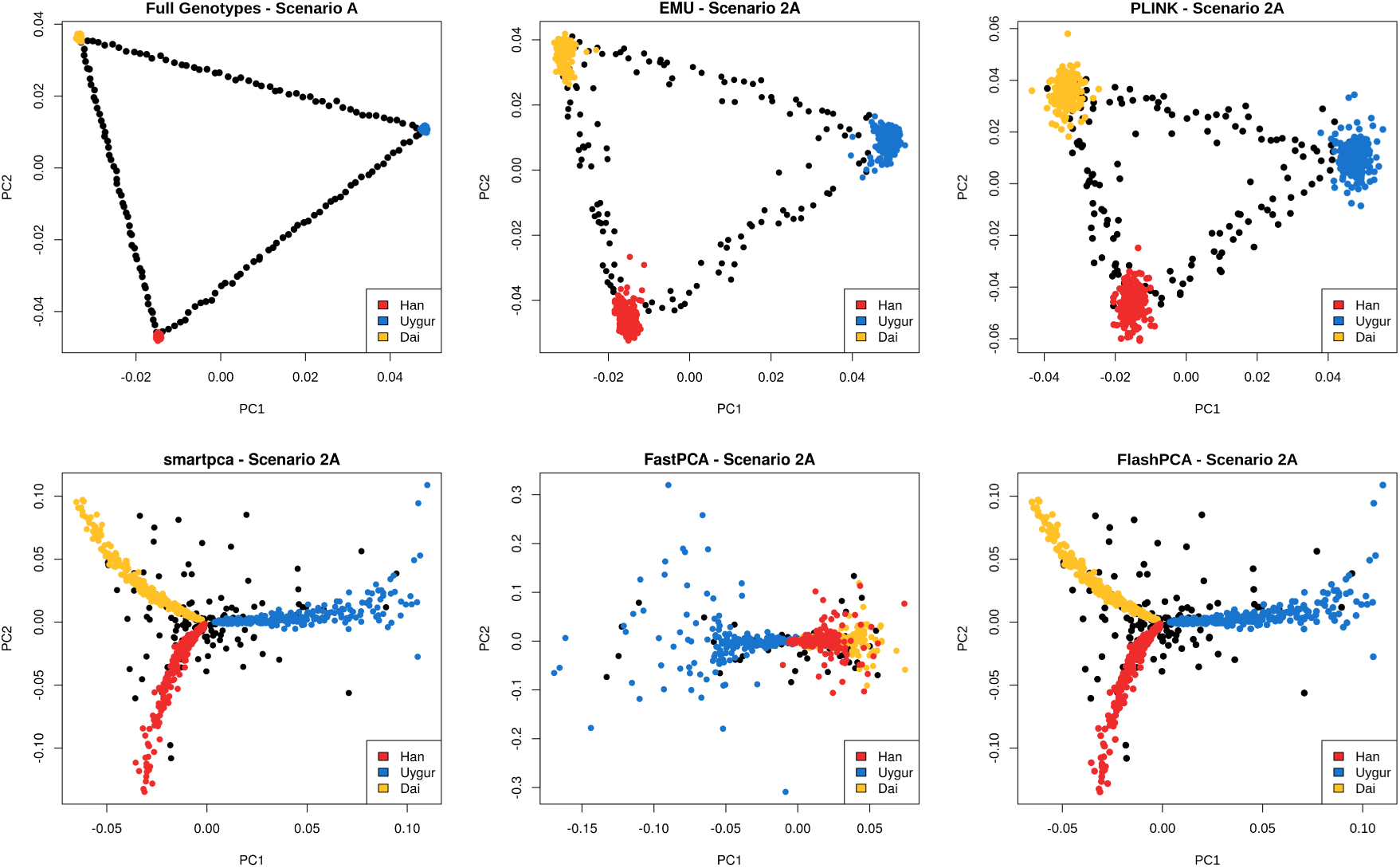
PCA plots of tested methods for Scenario 2A displaying the top two axes of genetic variation for 900 individuals. Black dots represent two-way admixed individuals. The top left plot shows the PCA performed on the full dataset, such that it acts as ground-truth. Individuals were simulated with extreme missingness rates between 90 − 99%.

In Scenario 3, the individuals sampled from the three population were simulated with two different settings of missingness and the results are displayed in Figure 3, S4 and S7. `EMU` and `PLINK` are capturing the population structure but `PLINK` still has more noisy estimates for the individuals of the three distinct populations. Due to the smaller variance in the missingness rate intervals, `smartpca`, `FastPCA` and `FlashPCA` are capturing the population structure accurately of the individuals simulated under one missingness setting. However, the individuals simulated under the other missingness setting now clusters together, which in particular illustrates the problem of not accounting for missingness as these clusters may be interpreted as separate populations.

**Figure 3:**
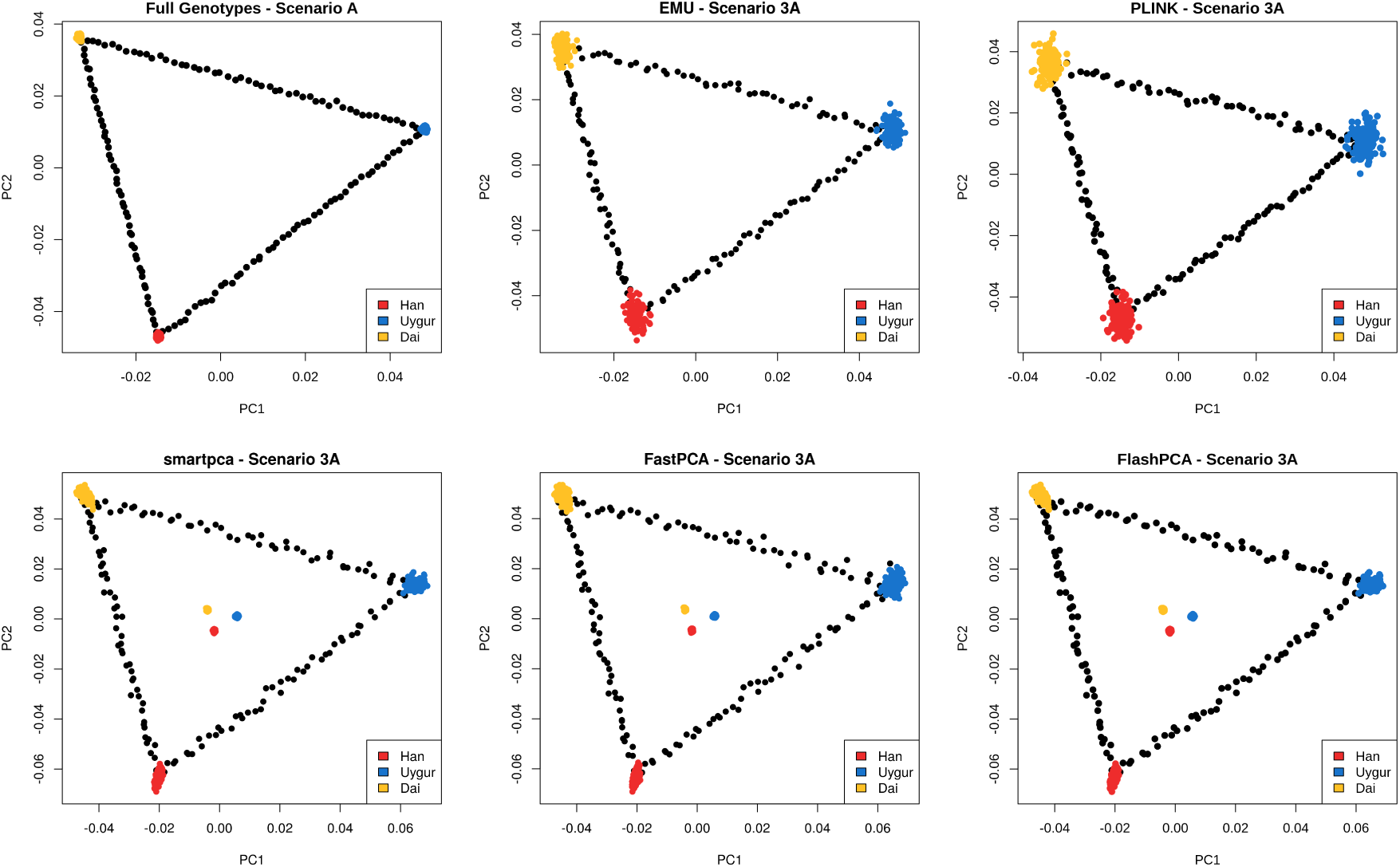
PCA plots of tested methods for Scenario 3A displaying the top two axes of genetic variation for 900 individuals. Black dots represent two-way admixed individuals. The top left plot shows the PCA performed on the full dataset, such that it acts as ground-truth. Individuals were simulated with missingness rates sampled as either ≈ 50% or ≈ 95%.

#### 3.1.2 SNP Ascertainment scheme scenarios

For the SNP ascertainment scenarios, we simulate 750 individuals from the three populations with 250 from each, thus excluding admixed individuals. We note that `PLINK` could not be run for Scenario 7 and 8 as it relies on pairwise estimates across all samples and can not run if there is no overlap in variants between a single pair of individuals. This would be a potential problem in its usage for ultra-low coverage sequencing data.

For Scenario 4, we are emulating the merging of whole-genome sequencing data and SNP array data by having half of the individuals in each of the three populations only have information in variants with a MAF ≥ 0.25. The results are visualized in Figure S10. Once again the methods performing mean imputation are biased by the different missingness rate, which is now created by having different SNP ascertainment schemes instead of being random, while `EMU` and `PLINK` are able to infer correct population structure. As the simulated missingness is non-random, we also see that a method like MDS is failing at capturing population structure (Figure S16).

In Scenario 5, half of the Han population is simulated such that it only had a small overlap with the other half (13K), while the two other populations are simulated with full information. The results are shown in Figure S11. All methods performing mean imputation interpret the two halves of the Han population as two separate populations, while `EMU` and `PLINK` still infer correct population structure.

Scenario 6 are similar to Scenario 5, but now two of the populations are simulated with two different halves (Han and Uygur). The results are visualized in Figure S12. Even with almost no overlap between the halves within two of the populations, `EMU` and `PLINK` are able to infer correct population structure. Of course, it is seen that mean imputation methods are biased once again and they show the four halves as different clusters.

The last two SNP ascertainment scheme scenarios are almost simulated identically as 5 and 6 but with no overlaps in the subsets of a population. We see almost the exact same results as for the previous two scenarios except that `PLINK` could not be run, and `EMU` is now failing to converge in the last scenario as there is no solution for the iterative procedure. The results are visualized in Figure S13 and S14 for Scenario 7 and 8, respectively.

### 3.2 Computational tests

We have tested runtimes and memory usage of `EMU` for different simulated sample sizes and compared it to the other tested methods. The different datasets were simulated under the same settings as Scenario 2 with individual missingness rates between 90−99%. The different sample sizes simulated were 900, 9000, 18000, 36000, 54000 and 90000. `smartpca` could not be run for sample sizes > 36000. All datasets have ∼350K variable sites after filtering out rare variants with a threshold of 5%.

All analyses were performed server-side using 64 threads (2.10 GHz; Intel Xeon Gold 6152), and the results of the computational tests are summarized in Figure 4. It is clear that the methods based on low-rank approximation (`EMU`, `FastPCA` and `FlashPCA`) are faster than the methods that construct the GRM followed by eigendecomposition. However, the implementation in `PLINK` does keep up with the low-rank approximation approaches to a certain extent but its approach would become unfeasible for large sample sizes. To demonstrate this, we also performed curve fitting of the runtimes for `EMU` and `PLINK` for comparison with more extreme sample sizes. The curves are shown in Figure S9. Here we can derive from extrapolation that it would take `PLINK` ∼98 days to infer population structure for 1 million individuals, while it would only take ∼4.1 hours for `EMU`. This is due to `PLINK` having to construct the GRM (𝒪(*mn*^2^)) and additionally having to perform eigendecomposition on it.

**Figure 4:**
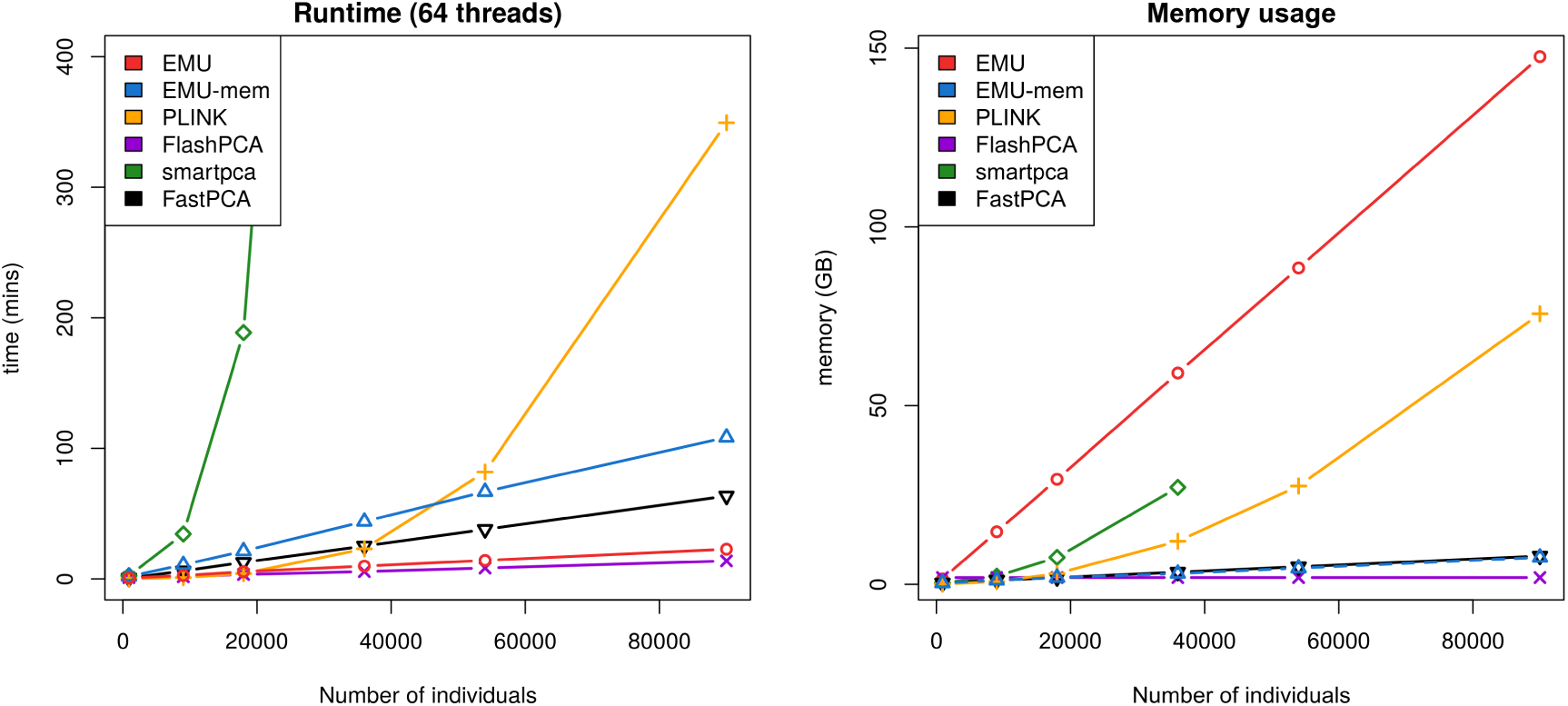
Computational tests. The left plot displays the average runtimes from 3 runs of the different methods for sample sizes of 900, 9000, 18000, 36000, 54000 and 90000. The number of sites is ∼350K for all simulated datasets. The right plot shows the memory usage of the different methods for the same datasets. `FlashPCA` used 2GB of memory for all evaluated sample sizes.

It is noteworthy that the number of iterations performed in `EMU` decrease when the sample size is increased (see Figure S8). This is due to having more individuals contributing to the axes of genetic variation, such that the eigenspectrum of the top principal components representing population structure is increased and random fluctuations are proportionally decreased. Thus, it becomes easier for `EMU` to impute the missing values in a more well-defined PC space.

### 3.3 Analysis on the 96,800 ultra low-pass genomes

We apply `EMU` and `PLINK` to analyse 96880 NIPT ultra-low pass genomes [21]. The participants came from 31 provinces throughout mainland China. After performing site filtering based on read length, sequencing error rate, minor allele frequency (≤ 0.05) and missingness rate (≤ 0.1), we obtain a genotype matrix consisting of 96,880 individuals and 440,183 sites. Again, both `EMU` and `PLINK` are run server-side with 64 threads under the same configurations as in the simulations. We use 3 eigenvectors to estimate individual allele frequencies in `EMU`. The inferred population structure of `EMU` and `PLINK` are visualized in Figure 5 and S15, respectively. Runtime information of both methods is displayed in Table 2, where `EMU` is shown to be ∼6.4x faster than `PLINK`. However due to the weak population structure in the Chinese individuals, `EMU` needs to perform 72 iterations that also naturally affects its runtime. As seen in the simulations, the results are similar but `EMU` seems to cluster the individuals better in comparison to `PLINK`. In the results, PC1 captures the cline of genetic variation from North to South China, while the PC2 captures the cline from West to East which is less apparent. PC2 is mainly driven by individuals with no province information. Some of these individual reported their ethnicity which were all Uygur – the main ethnic group in most western province (Xinjiang).

**Table 2:** Runtime and memory usage for dataset of 96,880 individuals and 440,813 sites.

|  | Runtime (mins) | Memory (GB) | Iterations |
| --- | --- | --- | --- |
| EMU | 75.2 | 198.8 | 72 |
| PLINK | 480.5 | 87.7 | N/A |

**Figure 5:**
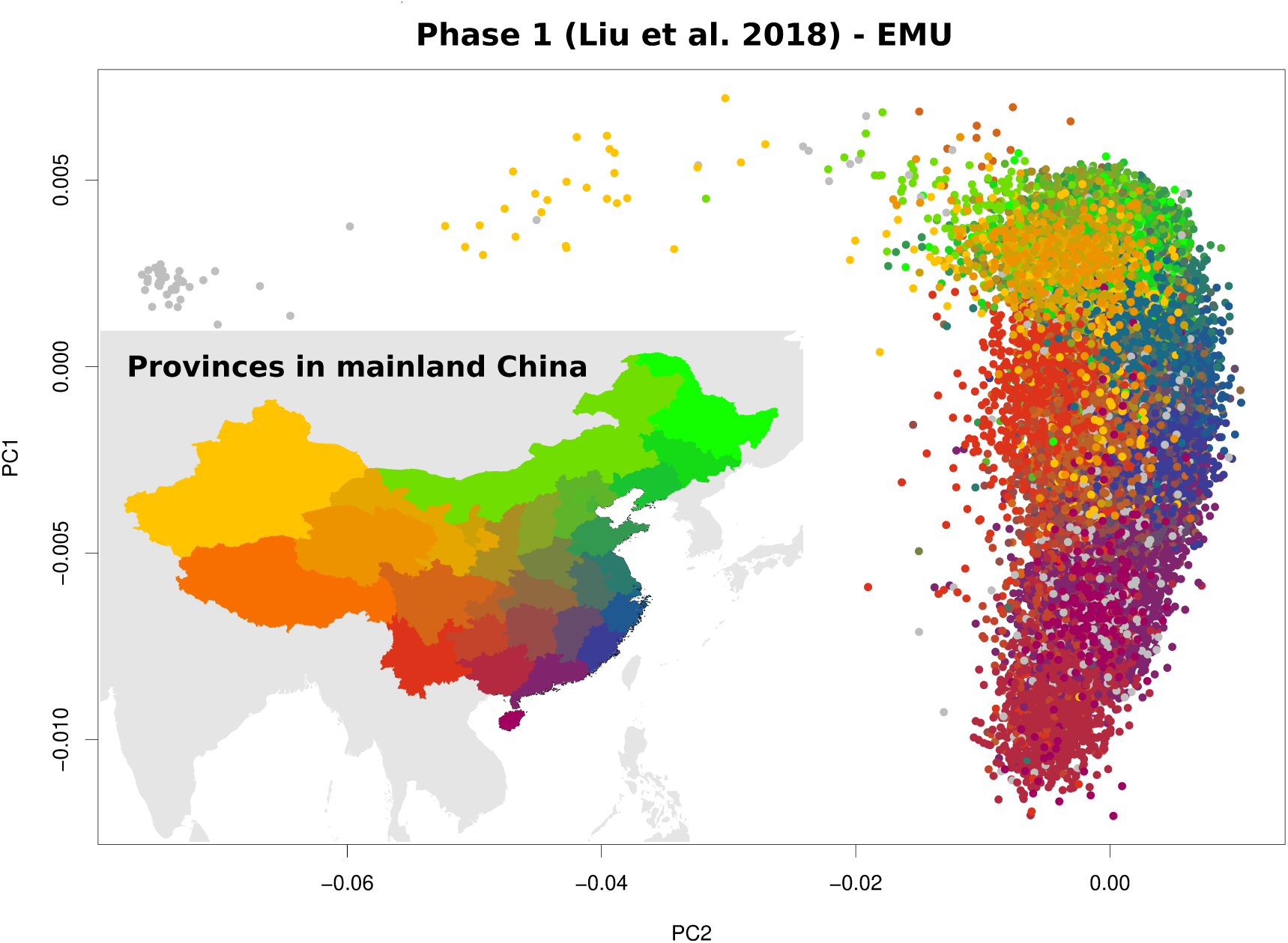
Inferred population structure using `EMU` on the Phase 1 dataset of 96,880 low-pass genomes. Individuals have been colored by their reported province of sampling where individuals colored grey have no information.

## 4 Discussion

We have implemented a novel method, `EMU`, for inferring population structure in large-scale genetic studies that accounts for random and non-random missingness. It estimates individual allele frequencies in an iterative manner based on low-rank approximations to impute missing information in the dataset through an EM-PCA approach. We have shown that `EMU` outperforms other commonly used approaches in population genetics and it is very competitive in terms of runtime for large-scale datasets.

Through simulation studies, we demonstrated that the majority of existing methods does not account for missingness in a meaningful way, which introduces major biases in the results of these methods. This bias is even observed for low missingness rates. `PLINK` also corrects for missingness and was shown to infer accurate population structure. However its approach is not feasible for very large-scale genetic studies as it performs full eigendecomposition on the constructed GRM. By extrapolation from runtimes of computational tests, `PLINK` would need to run for ∼98 days (compared to ∼4.1 hours for `EMU`) to infer population structure for 1 million individuals under the tested configurations of this study. We further note that `EMU` outperformed `PLINK` in every scenario regarding inference of principal components.

As previously mentioned, the results of `smartpca`, `FlashPCA` and `FastPCA` were almost seen to be identical for all the tested scenarios, however `FastPCA` deviated from the other two in Scenario 2A, 2B and 3A where it performed slightly worse. This is due to `FastPCA`, also based on random matrix theory, not performing any power iterations including normalizations for stabilizing results as done in the randomized PCA procedure [14] (`EMU`) as well as methods based on `ARPACK` [19] procedures (`FlashPCA`). This shows the importance of having stabilization steps in small noisy datasets, as the results of `FlashPCA` are similar to `smartpca` that performs full eigendecomposition on the GRM with mean imputation, and thus appear more robust.

We also applied `EMU` to the dataset of Phase 1 study of the Chinese Millionome Project, where we show that `EMU` is able to reconstruct the previous findings in a matter of CPU hours compared to CPU hours of the original study. We are able to identify population structure across mainland China with two clines from North to South and from West to East, respectively.

One limitation of the iterative nature of `EMU` is the fact that the individual allele frequencies need to be updated, and thus stored in memory. This can become a problem for large-scale genetic studies with sample sizes greater than 100K to run on standard server solutions. We therefore also implemented a more memory-efficient variant of `EMU`, which only keep the low-rank factor matrices in memory and compute individual allele frequencies when needed at the cost of computational speed. We also believe that our method may be of use in the field of applied statistics as we successfully combine EM-PCA with a SQUAREM acceleration scheme on top of truncated SVD methods to infer population structure in the presence of large missingness.

## 4.1 Acknowledgements

The study was supported by the Lundbeck foundation (R215-2015-4174) (A.A & J.M.). S.L. & M.H. was supported by National Natural Science Foundation of China (31900487).

## 4.2 Author contributions

J.M. & A.A. developed the method. J.M. wrote the algorithm, wrote the manuscript and performed simulation analyses. S.L. & M.H. performed the analysis using the NIPT data. All authors reviewed and contributed to the final manuscript.

## Supplementary Material

### EMU-mem **- Memory-efficient implementation**

Here we describe a more memory-efficient implementation of `EMU`. Instead of computing and storing the individual allele frequencies 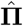 after performing SVD (step 3, algorithm 1), then only the decomposition matrices 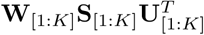 are retained at each iteration. By only using these matrices alongside the original data matrix **D** and population allele frequencies 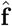, we will never have to construct the memory heavy matrix **E** to keep track of the individual allele frequencies. We will instead estimate the values that are needed on the fly. This is possible through the use of the low-rank approximation variant of Halko et al. [14], such that the decomposition of the truncated SVD is based on lazy evaluation of **E** in its matrix multiplication steps. Additionally, **D** is also stored in a 2-bit integer format as only 2-bits are needed for an individual in a given site. Thus each byte will contain the genotypes of 4 individuals.

Hence only **W**_[1:*K*]_, **S**_[1:*K*]_ and 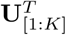 will be updated in each iteration.

### Memory usage of EMU variants

The memory usage in `EMU` is mainly governed by:

- **D** *n* × *m* `signed char` matrix,
- **E** *n* × *m* `float` matrix.

Such that roughly 5*nm* bytes are used in memory.

While for `EMU-mem`, the memory usage is mainly governed by:

- **D** *b* × *m* `unsigned char` matrix, where 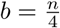.

This is a 95% reduction in memory usage compared to `EMU`, by using roughly 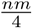 bytes.

## Supplementary Figures

**Figure S1:**
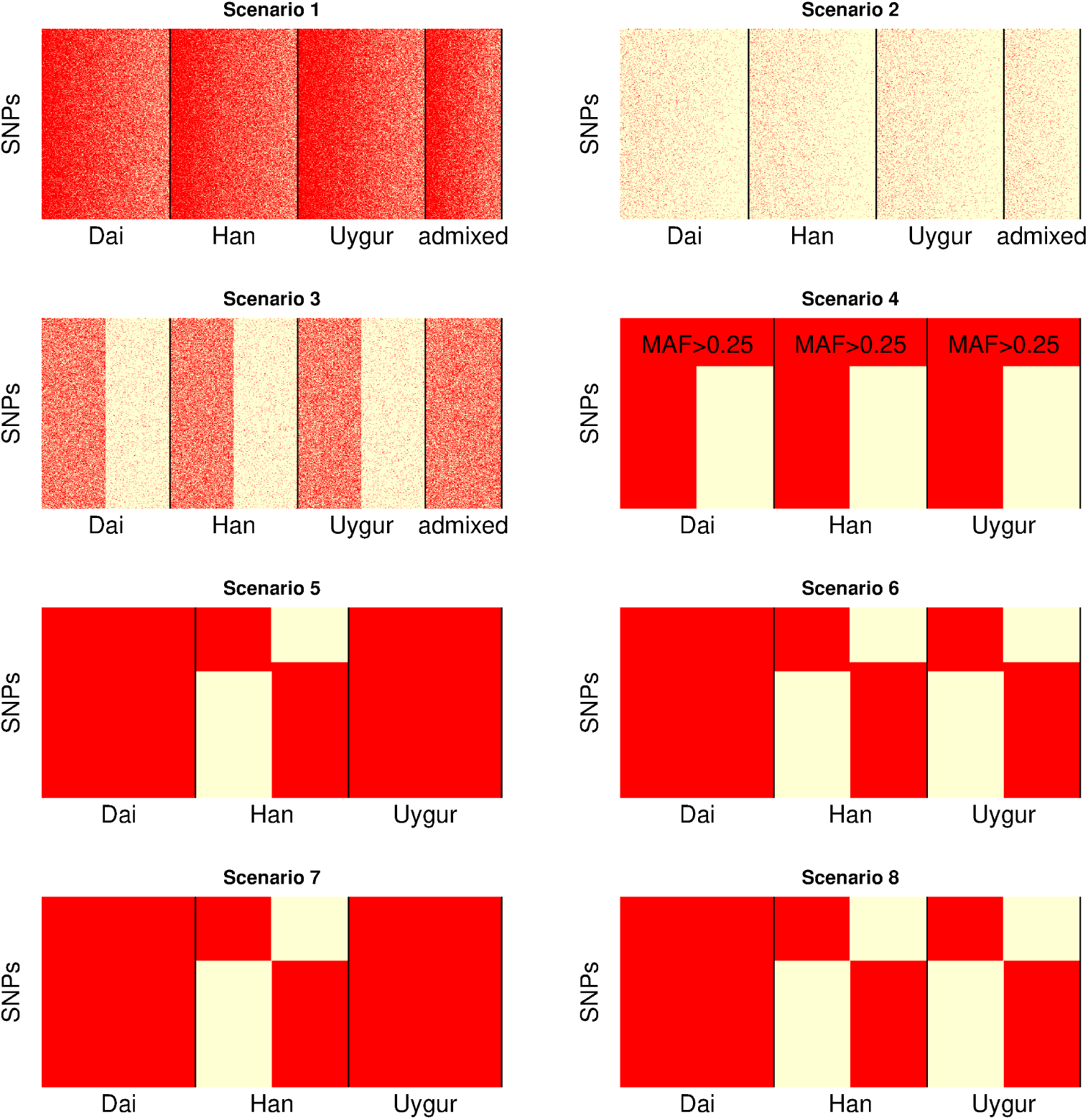
Graphical overview of the simulated scenarios. Rows represent SNPs and columns represent individuals and each dot is colored by missingness such that red means information while sand-white means missing value.

**Figure S2:**
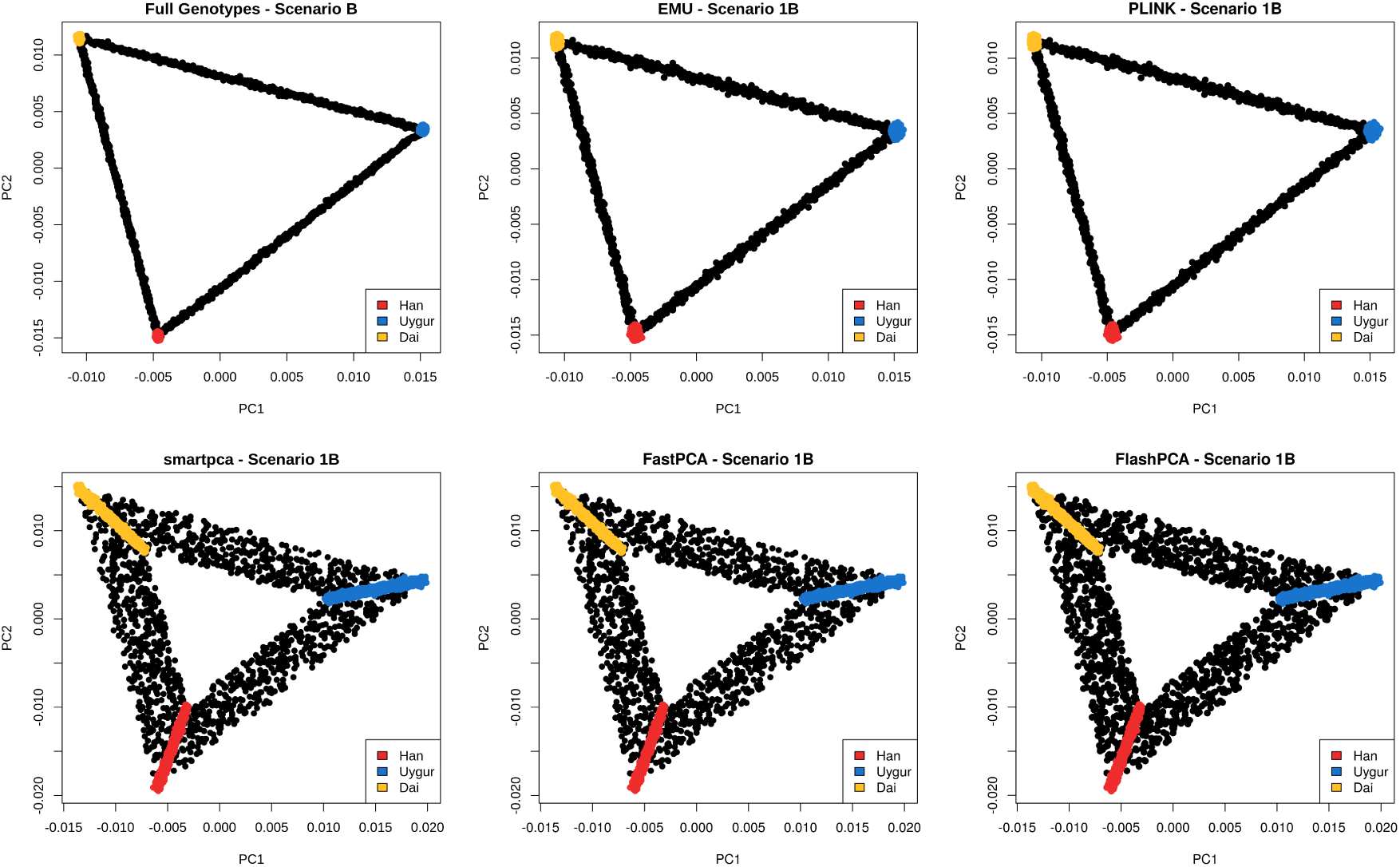
PCA plots of tested methods for Scenario 1B displaying the top two axes of genetic variation, similarly to Figure 1, however performed on 9000 individuals. Black dots represent two-way admixed individuals. The top left plot shows the PCA performed on the full dataset, such that it acts as ground-truth. Individuals were simulated with low to moderate missingness rates between 5 − 50%.

**Figure S3:**
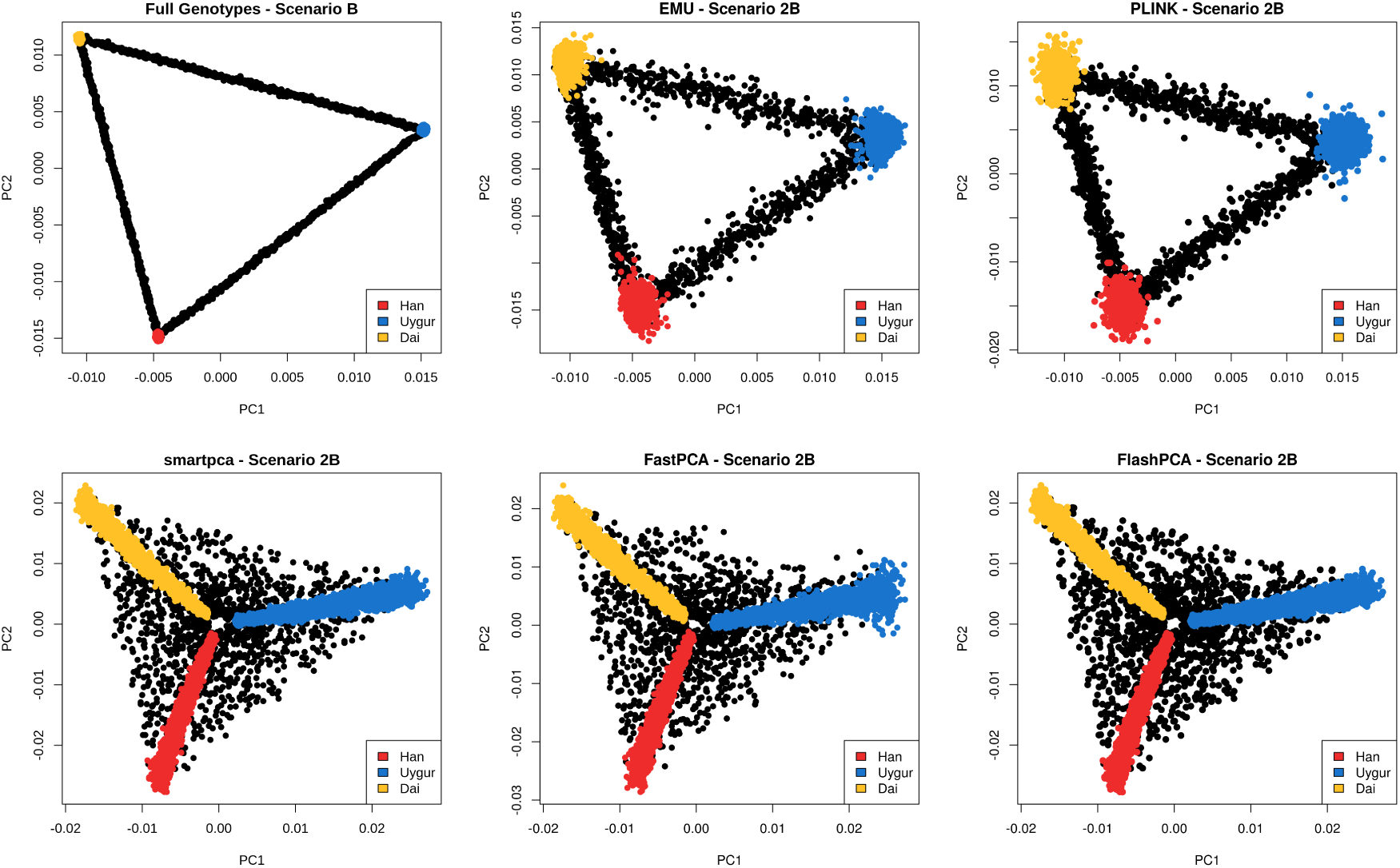
PCA plots of tested methods for Scenario 2B displaying the top two axes of genetic variation, similarly to Figure 2, however performed on 9000 individuals. Black dots represent two-way admixed individuals. The top left plot shows the PCA performed on the full dataset, such that it acts as ground-truth. Individuals were simulated with extreme missingness rates between 90 − 99%.

**Figure S4:**
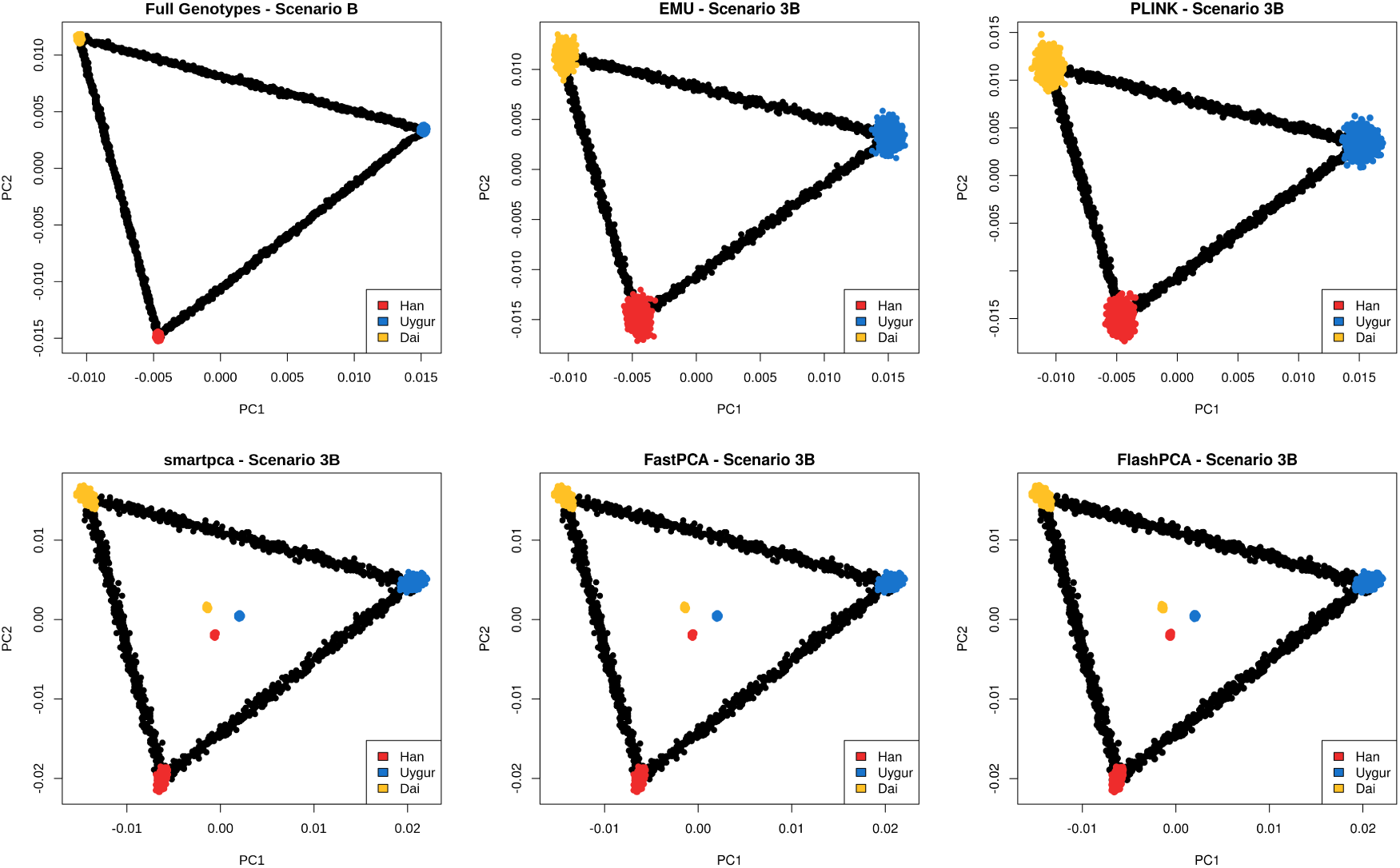
PCA plots of tested methods for Scenario 3B displaying the top two axes of genetic variation, similarly to Figure 3, however performed on 9000 individuals. Black dots represent two-way admixed individuals. The top left plot shows the PCA performed on the full dataset, such that it acts as ground-truth. Individuals were simulated with missingness rates sampled as either ≈ 50% or ≈ 95%.

**Figure S5:**
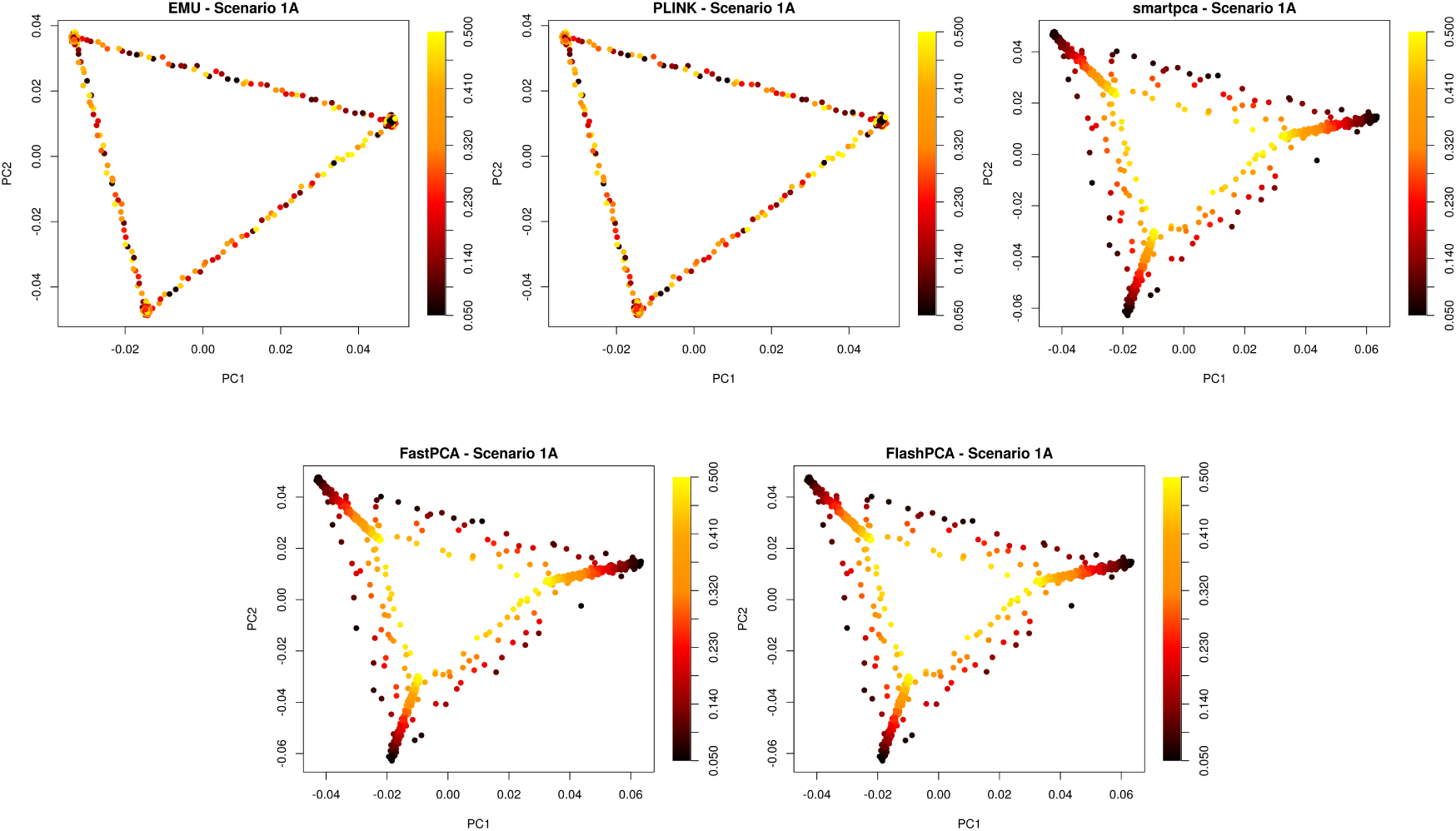
The same PCA plots as in Figure 1, however the individuals have been coloured by their sampled missingness rate. Here in Scenario 1A, the individuals were assigned rates between 5 − 50%.

**Figure S6:**
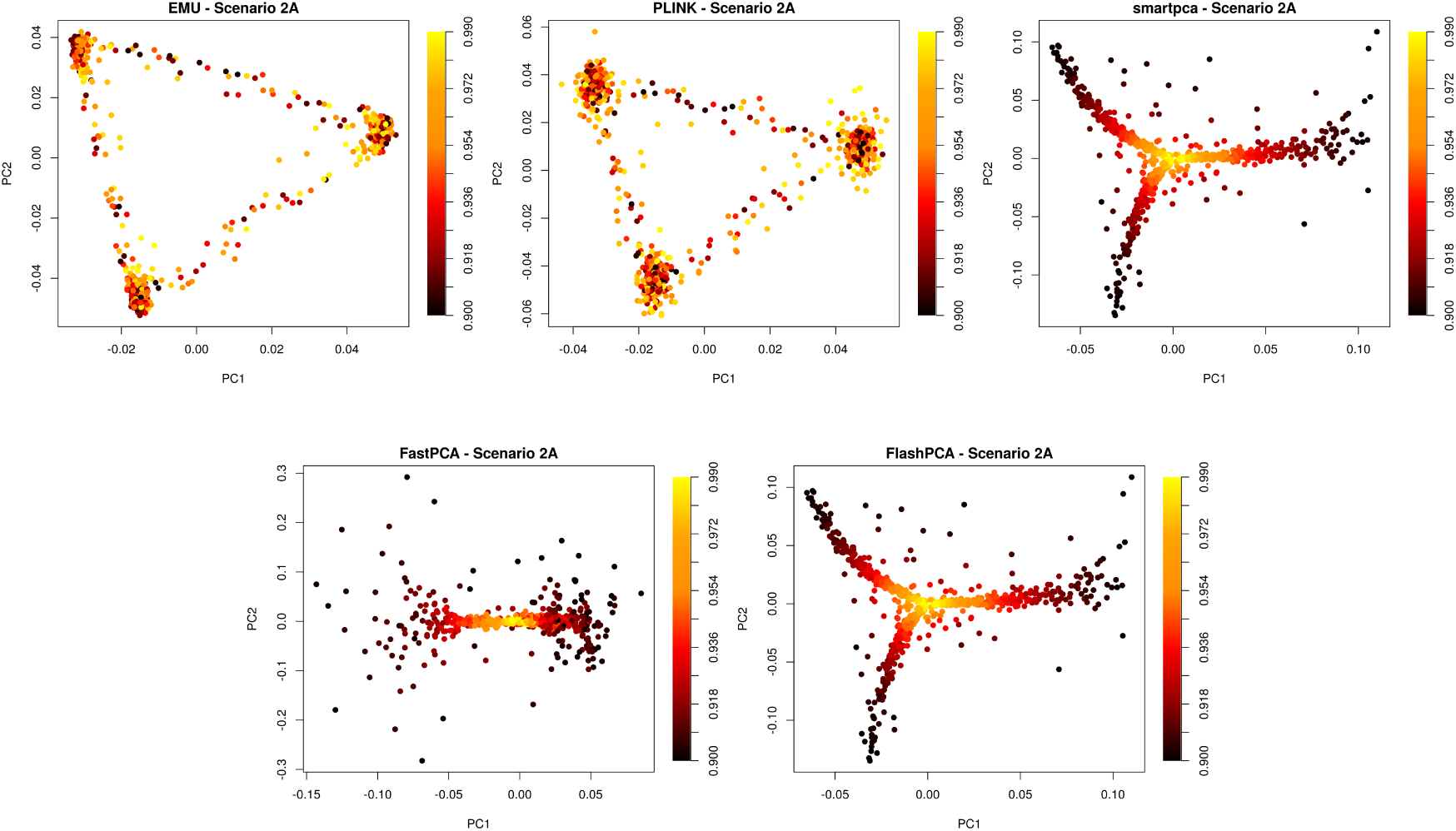
The same PCA plots as in Figure 2, however the individuals have been coloured by their sampled missingness rate. Here in Scenario 2A, the individuals were assigned rates between 90 − 99%.

**Figure S7:**
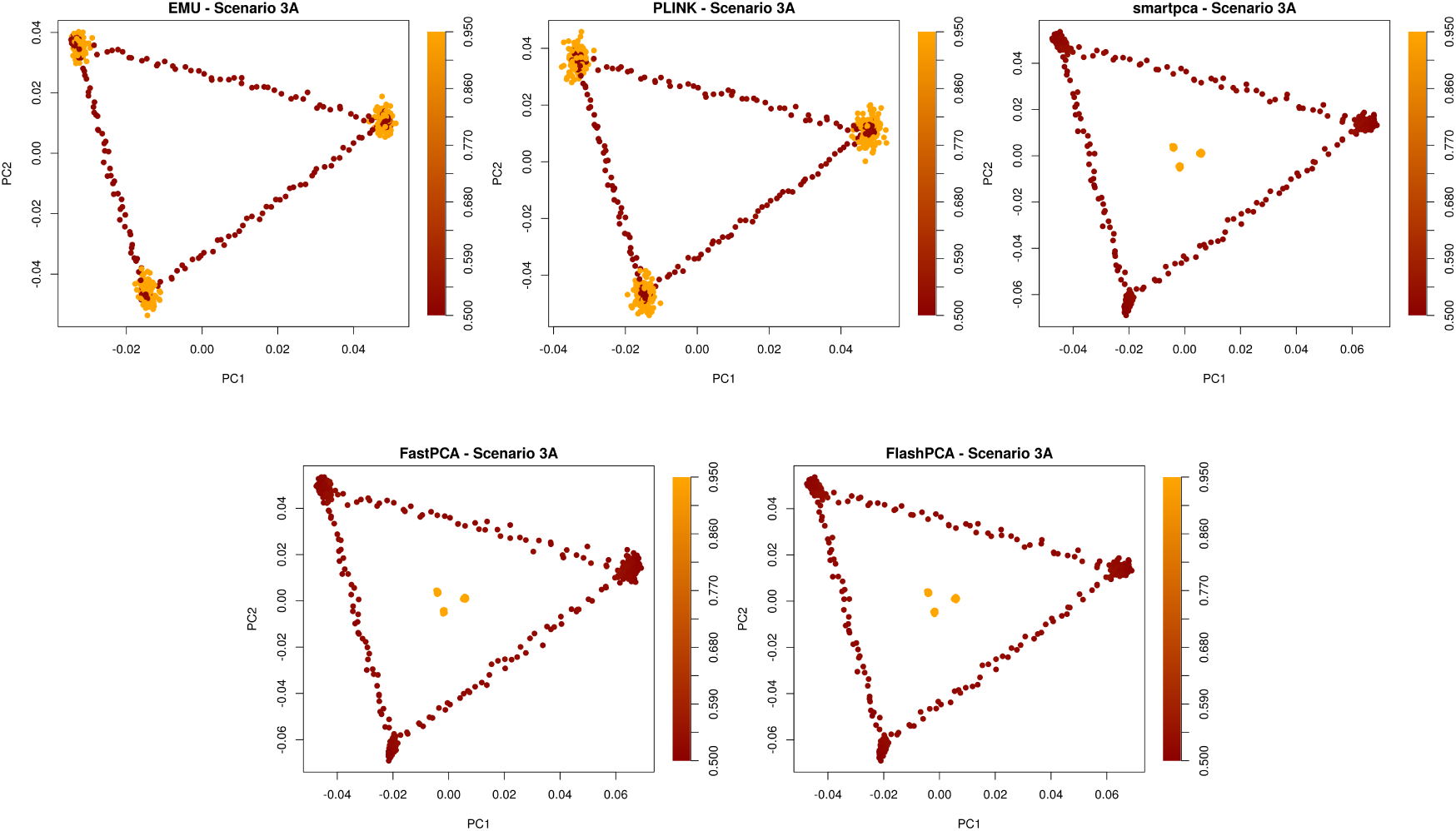
The same PCA plots as in Figure 3, however the individuals have been coloured by their sampled missingness rate. Here in Scenario 3A, the individuals were assigned rates around either 50% and 95%.

**Figure S8:**
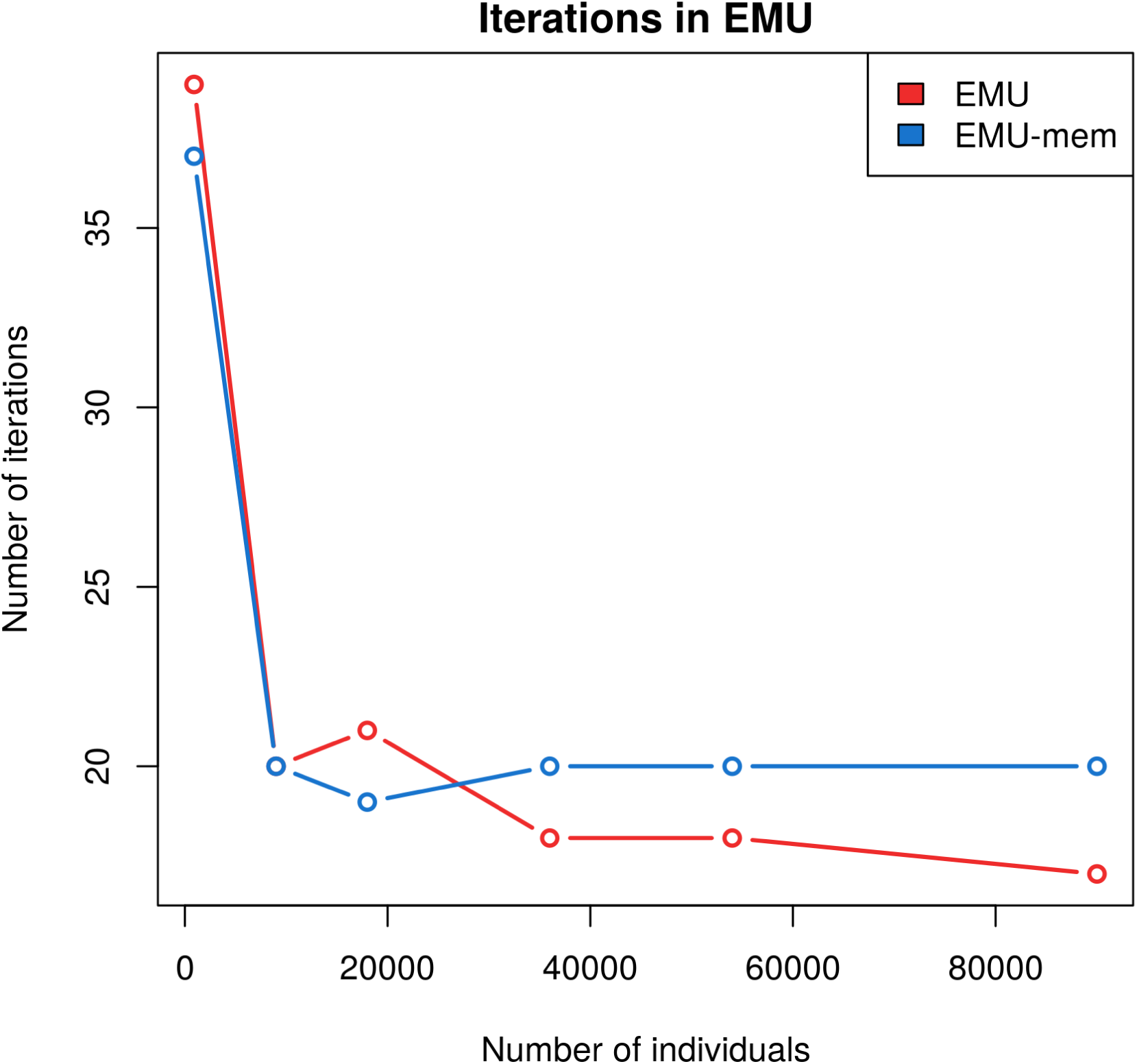
Number of iterations until convergence in `EMU` and `EMU-mem` as a function of the sample size of the dataset.

**Figure S9:**
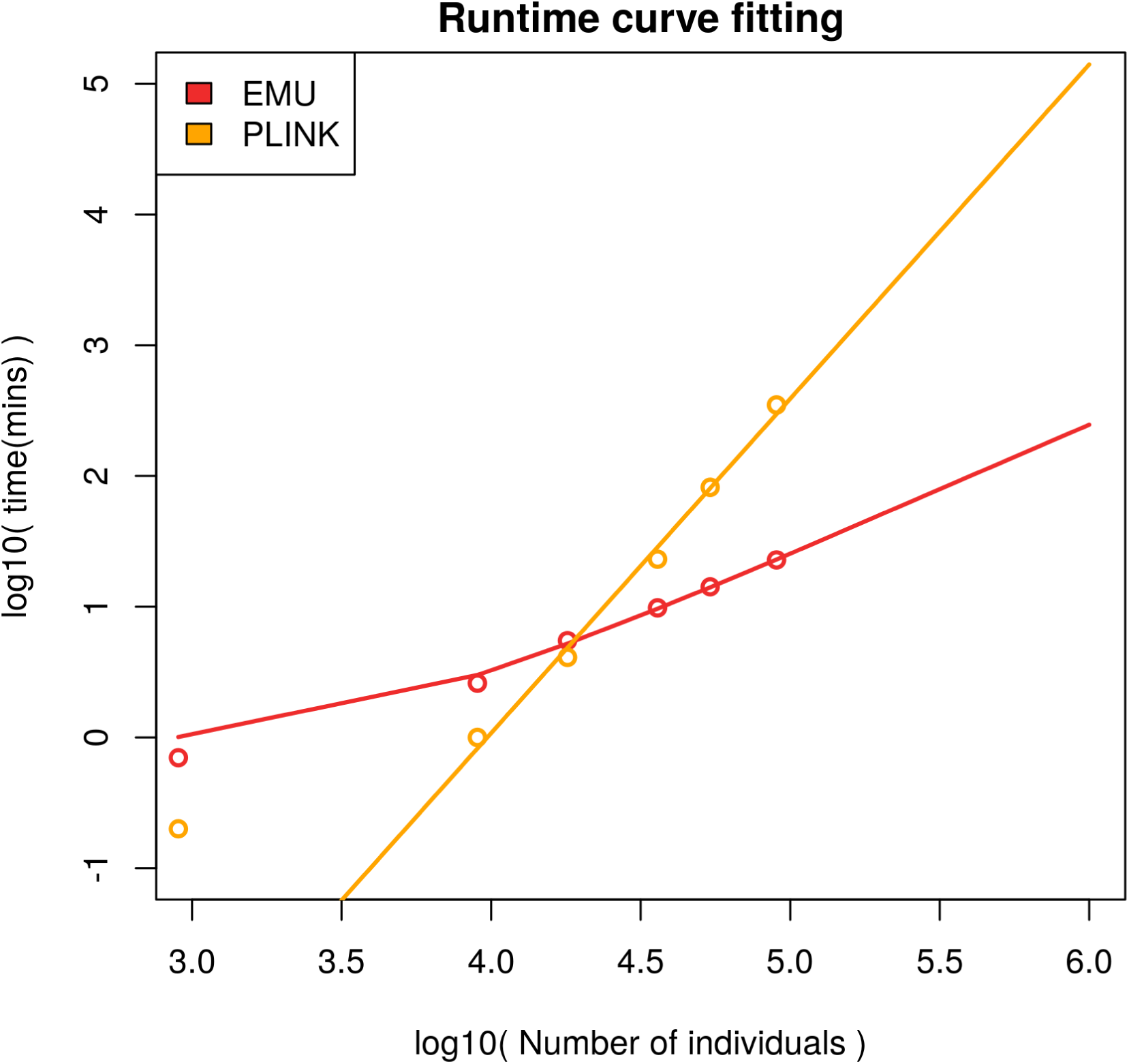
Curve fitting of runtimes for `EMU` and `PLINK`. Data points and fitted lines are in log_10_ scaling. A linear model has been fitted for `EMU` and an exponential model for `PLINK`. Assuming that the number of iterations of `EMU` will not decrease further, then the runtime can be approximated by the following function: *f*(*n*) = 0.000246*n* + 0.786. While the runtime for PLINK is approximated by: *g*(*n*) = 10^−10.2^ × *n*^2.558^. Here *n* is the number of individuals, while the number of sites is assumed constant (350K). The points represent the actual runtimes, while the lines follow the fitted runtimes. The first runtime point was not included in the curve fitting.

**Figure S10:**
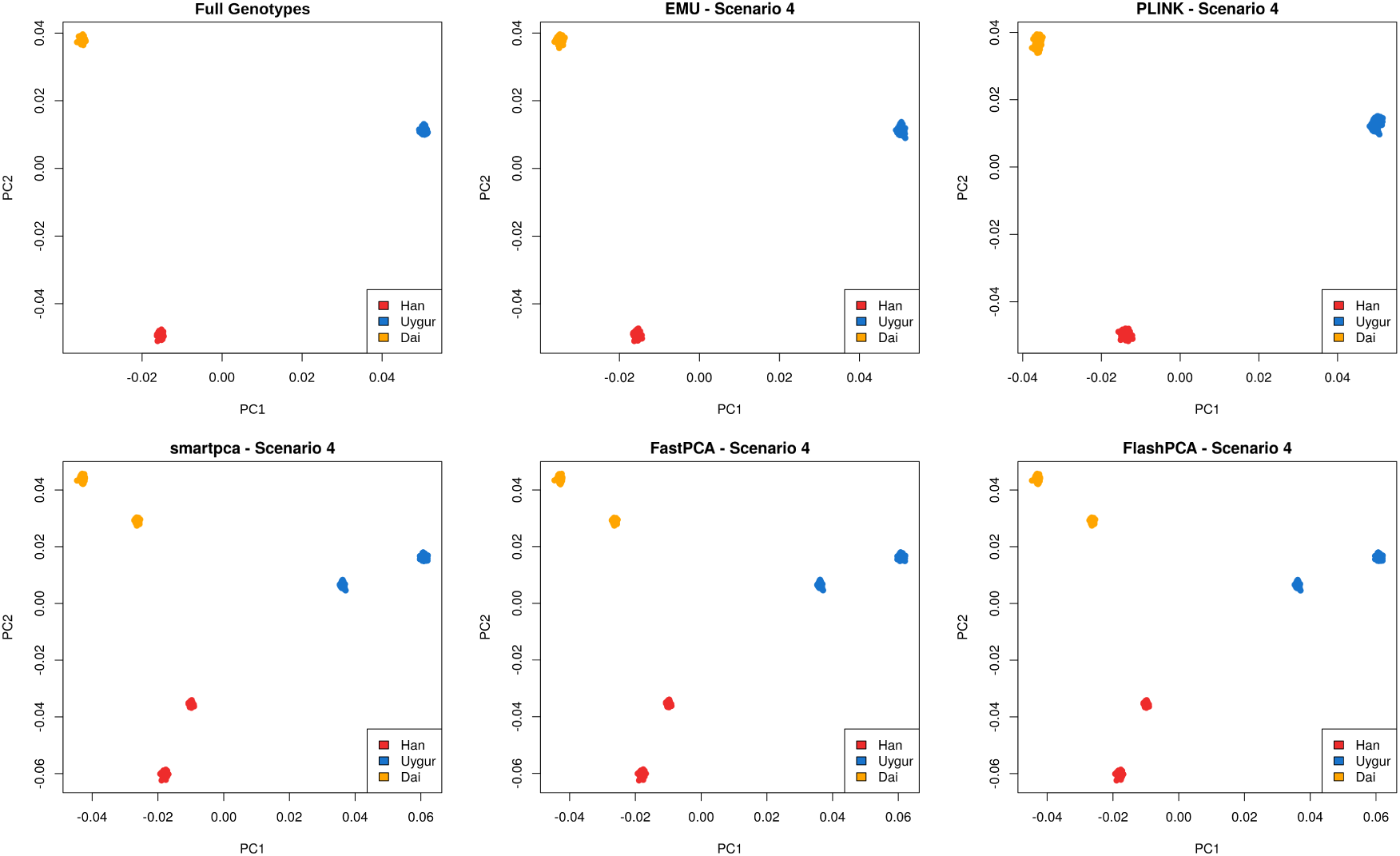
PCA plots of tested methods for Scenario 4 displaying the top two axes of genetic variation. The top left plot shows the PCA performed on the full dataset, such that it acts as ground-truth. Half of the individuals from each population were simulated using only common variants (MAF ≥ 0.25), while the of the halves were simulated using the entire set of variants to emulate the merging of datasets from SNP arrays with whole-genome sequencing.

**Figure S11:**
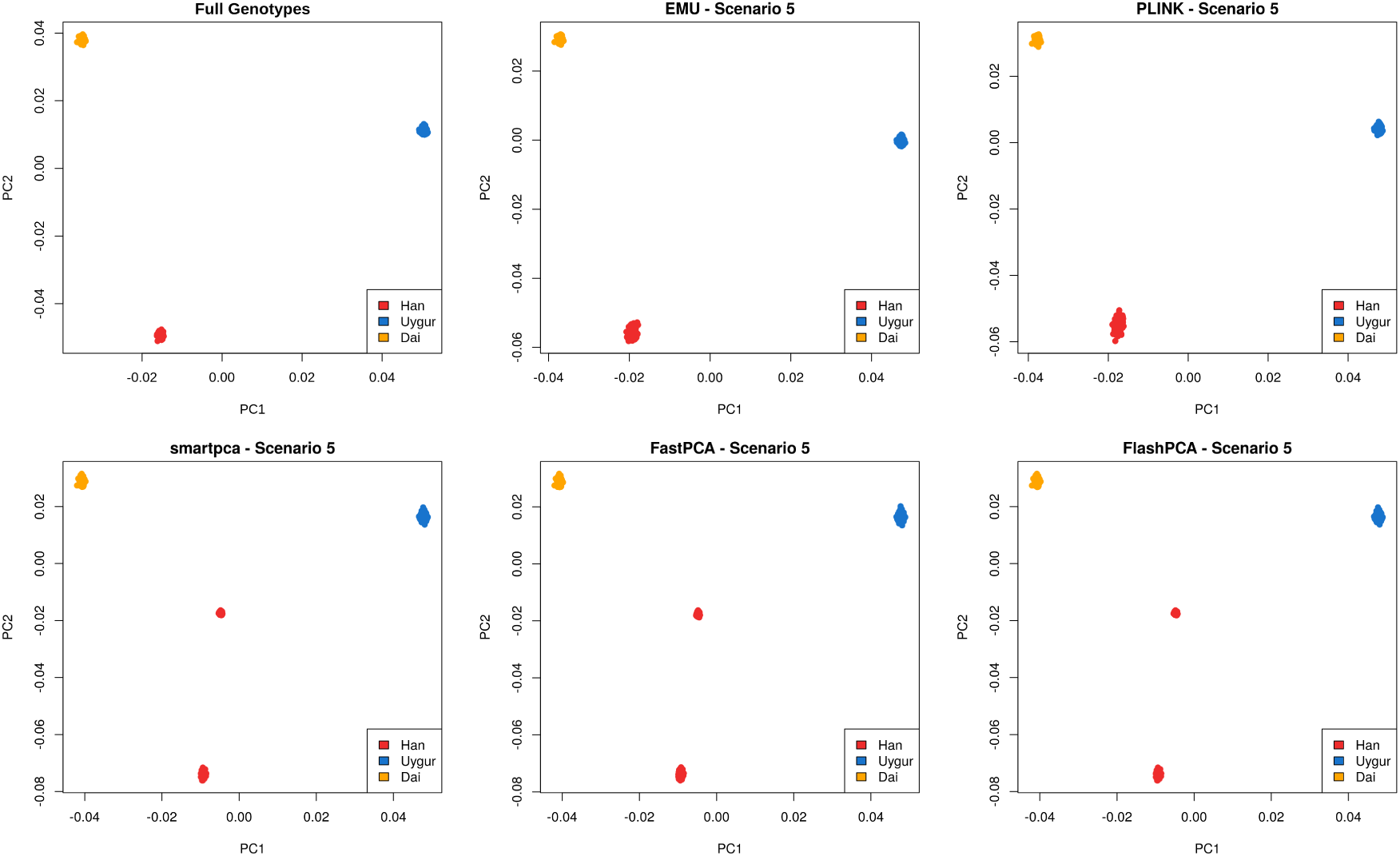
PCA plots of tested methods for Scenario 5 displaying the top two axes of genetic variation. The top left plot shows the PCA performed on the full dataset, such that it acts as ground-truth. In this scenario, one half of the simulated Han individuals only has information in ∼130K variants, while the other half has information in ∼233K variants with ∼13K variants overlapping between the halves. The other two populations have full information.

**Figure S12:**
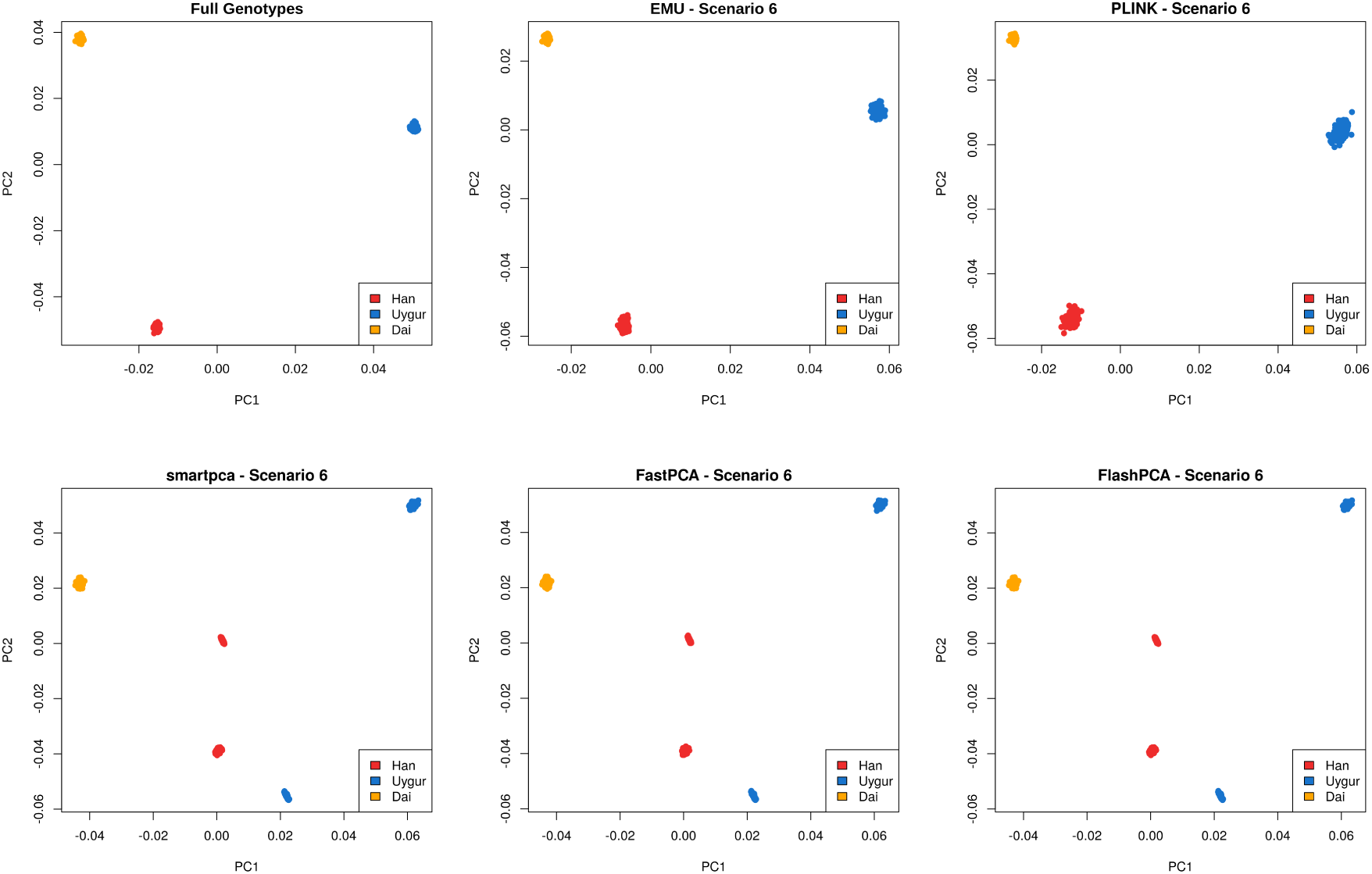
PCA plots of tested methods for Scenario 6 displaying the top two axes of genetic variation. The top left plot shows the PCA performed on the full dataset, such that it acts as ground-truth. The simulation procedure is almost identical to Scenario 5 but it is performed for two populations (Han and Uygur) instead of one. The Dai population is simulated with full information.

**Figure S13:**
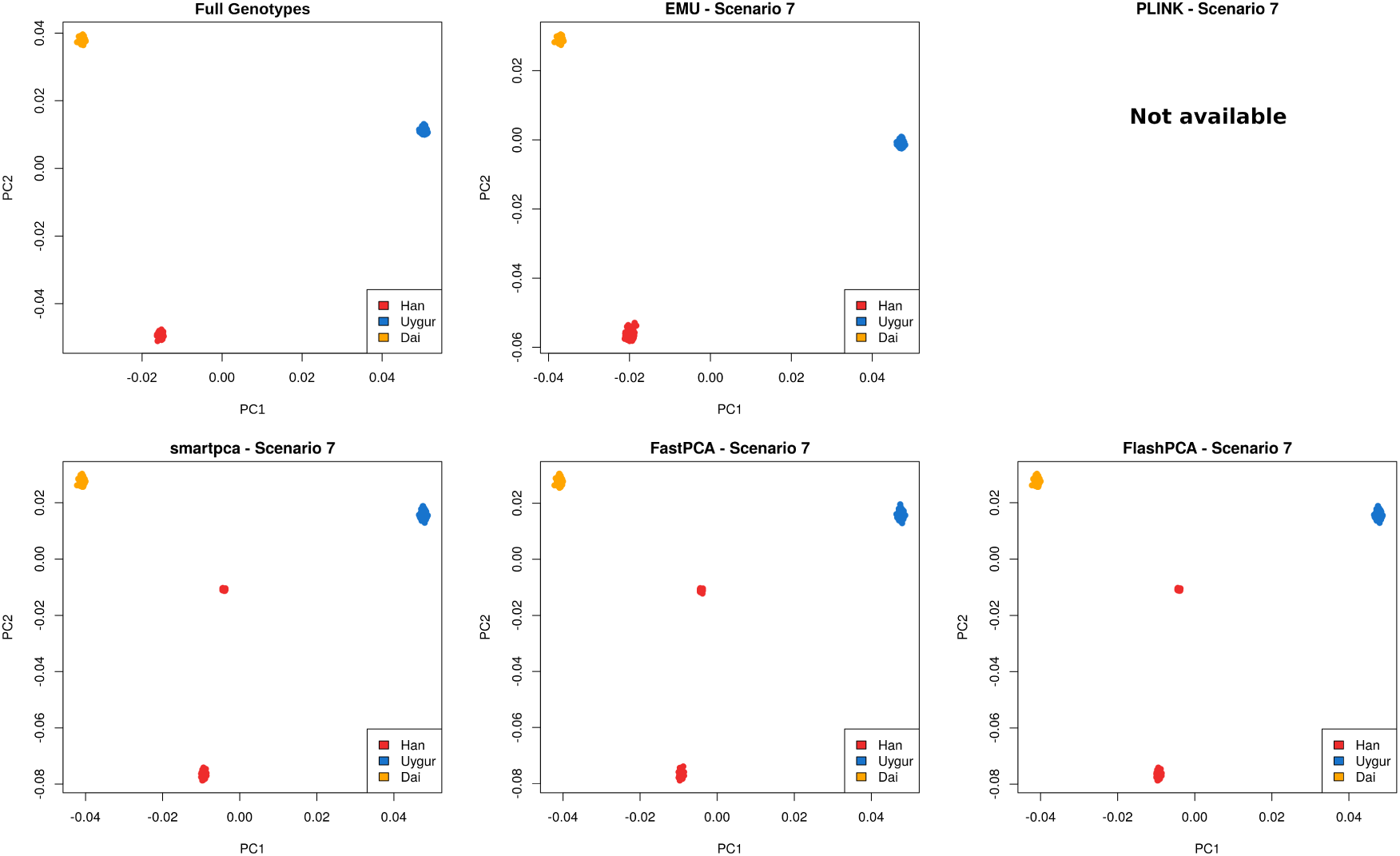
PCA plots of tested methods for Scenario 7 displaying the top two axes of genetic variation. The top left plot shows the PCA performed on the full dataset, such that it acts as ground-truth. The simulation scenario is almost identical to Scenario 5, however there is no overlap in variants between the two subsets of the Han population. Thus, one half of the simulated Han individuals only has information in ∼117K variants while the other half has information in ∼233K variants. `PLINK` was not able to run in this scenario.

**Figure S14:**
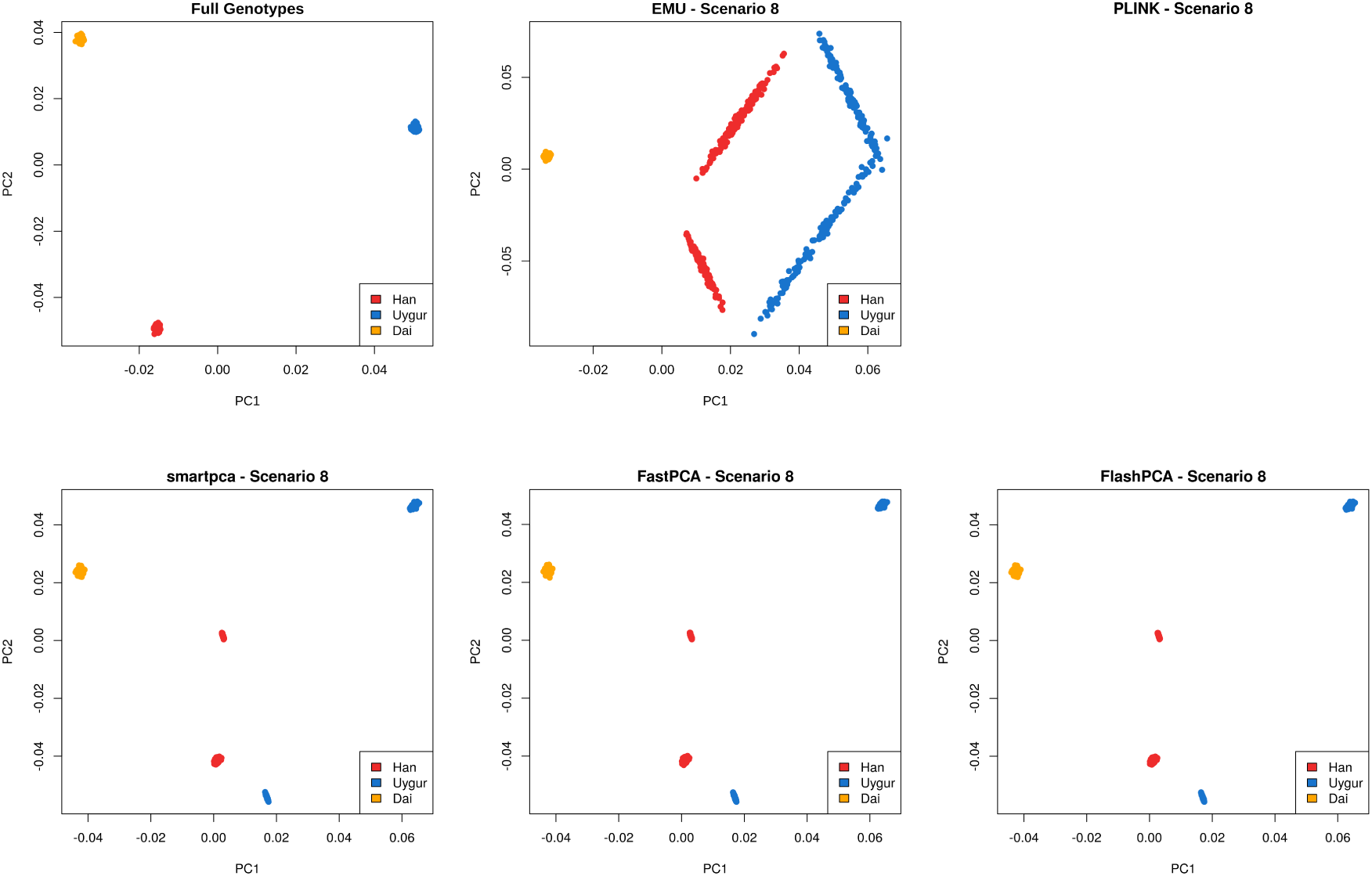
PCA plots of tested methods for Scenario 8 displaying the top two axes of genetic variation. The top left plot shows the PCA performed on the full dataset, such that it acts as ground-truth. The simulation procedure is almost identical to Scenario 7 but it is performed for two populations (Han and Uygur) instead of one. The Dai population is simulated with full information. `PLINK` was not able to run in this scenario.

**Figure S15:**
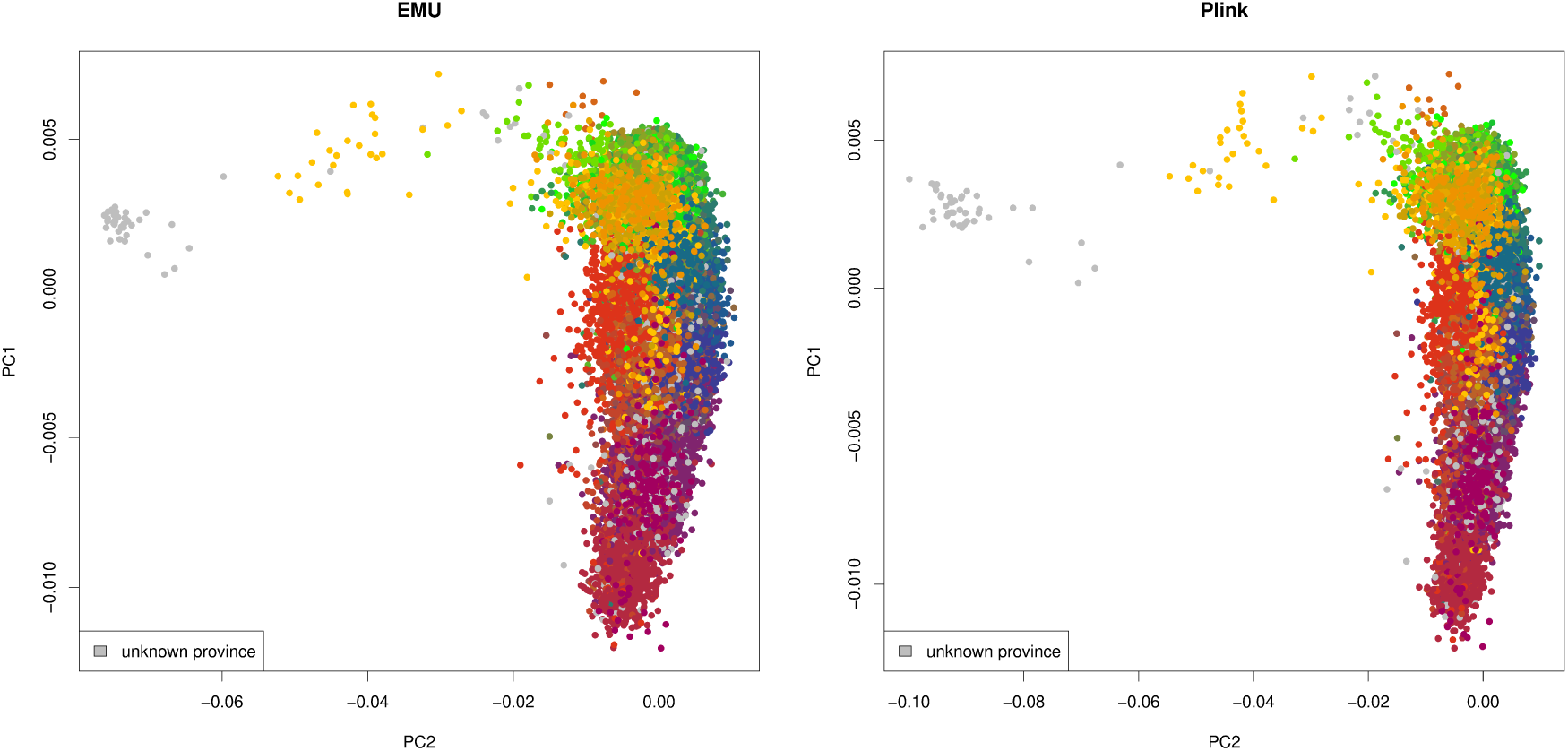
Inferred population structure using `EMU` and `PLINK` on the Phase 1 dataset of 96,880 low-pass genomes. Individuals have been colored by their reported province of sampling where individuals colored grey have no information..

**Figure S16:**
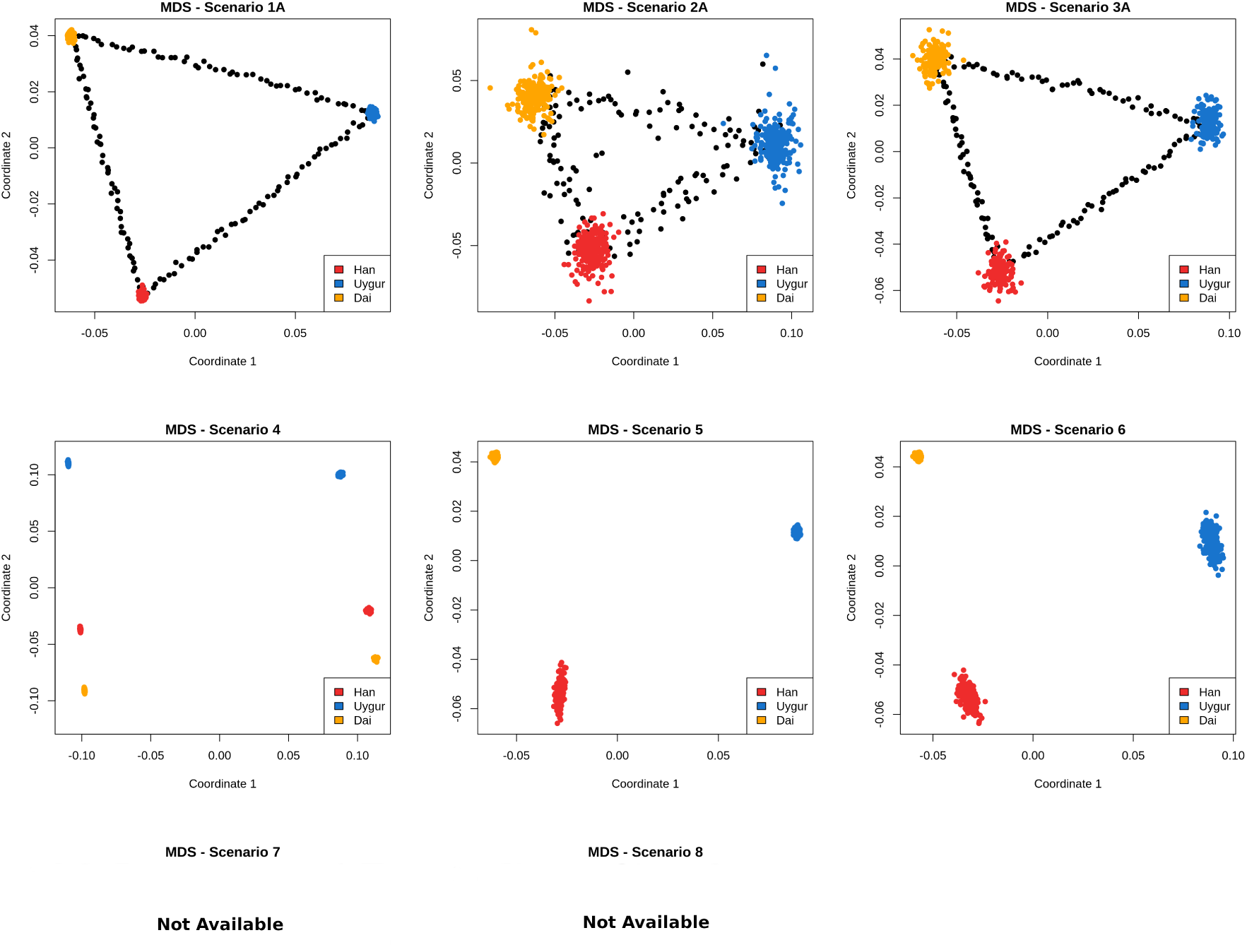
MDS using `PLINK` for all simulated scenarios. Black dots represent two-way admixed individuals. MDS could not be run for the last two scenarios due to the same reason as for `PLINK`.

**Figure S17:**
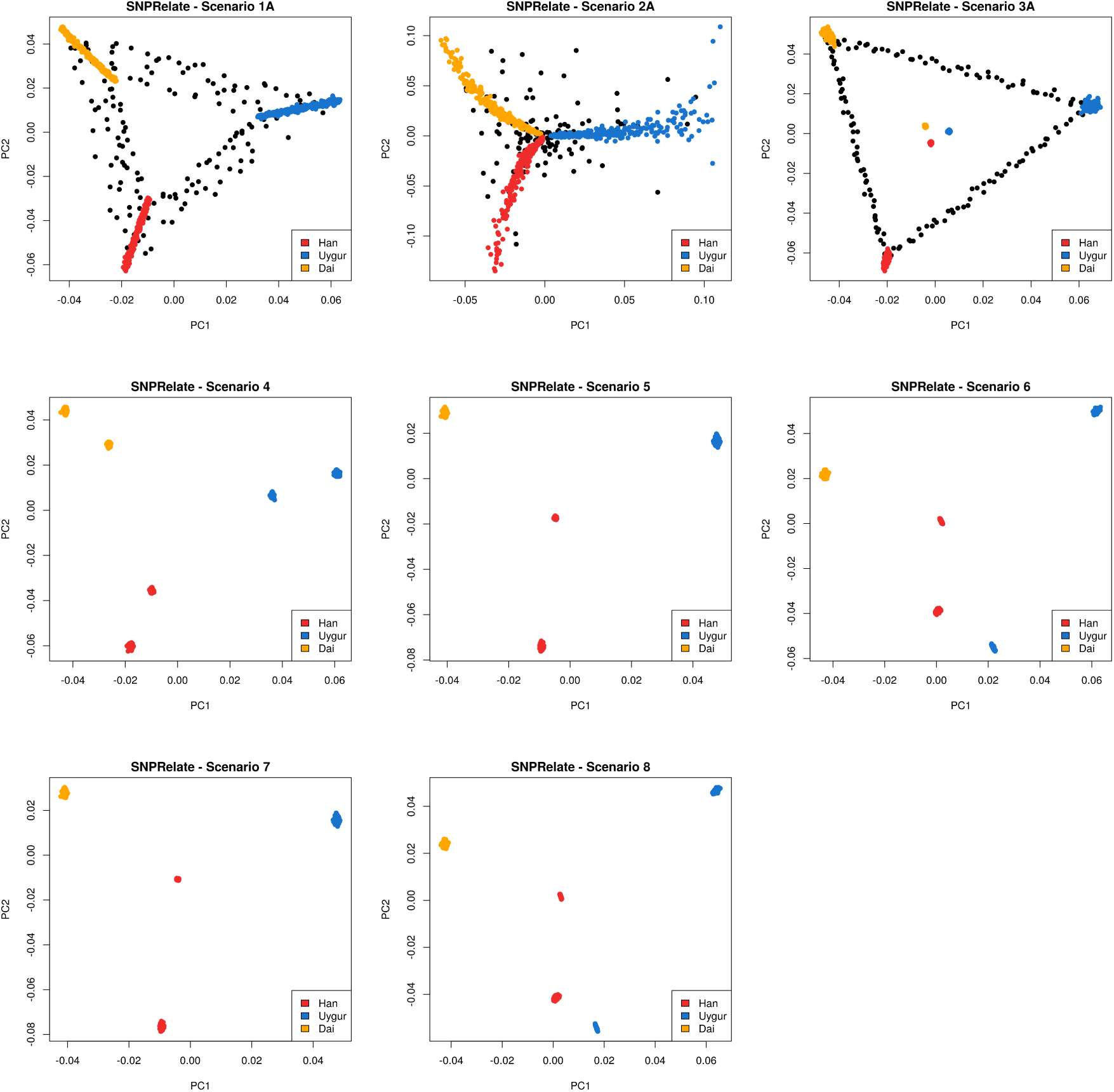
PCA plots using the `R` package `SNPRelate` for all simulated scenarios. `SNPRelate` suffers from the bias of mean imputation as seen in the other methods. Black dots represent two-way admixed individuals.

